# Myeloid mechano-metabolic programming restricts anti-tumor immunity

**DOI:** 10.1101/2022.07.14.499764

**Authors:** K.M. Tharp, K. Kersten, O.M. Maller, G.A. Timblin, C. Stashko, F.P. Canale, M-K. Hayward, I. Berestjuk, J. ten Hoeve-Scott, B. Samad, A.J. Ironside, R. Geiger, A.J. Combes, V.M. Weaver

**Affiliations:** Center for Bioengineering and Tissue Regeneration, Department of Surgery, University of California San Francisco, San Francisco, CA 94143, USA; Department of Pathology, University of California San Francisco, San Francisco, CA 94143, USA; ImmunoX Initiative, University of California San Francisco, San Francisco, CA 94143, USA; Institute for Research in Biomedicine, Università della Svizzera italiana, Bellinzona, Switzerland; UCLA Metabolomics Center, Department of Molecular and Medical Pharmacology, University of California, Los Angeles, Los Angeles, CA 90095, USA; UCSF CoLabs, University of California San Francisco, San Francisco, CA 94143, USA; Department of Pathology, Western General Hospital, NHS Lothian, Edinburgh, UK; Department of Medicine, Gastroenterology division, University of California San Francisco, San Francisco, CA 94143, USA; Department of Bioengineering and Therapeutic Sciences and Department of Radiation Oncology, Eli and Edythe Broad Center of Regeneration Medicine and Stem Cell Research, and The Helen Diller Family Comprehensive Cancer Center, University of California San Francisco, San Francisco, CA 94143, USA

## Abstract

Tumor progression is accompanied by fibrosis, which is associated with diminished anti-tumor immune infiltrate. Here, we demonstrate that tumor infiltrating myeloid cells respond to the stiffened fibrotic tumor microenvironment (TME) by initiating a TGF-beta (TGFβ)-directed, collagen biosynthesis program. A collateral effect of this programming is an untenable metabolic milieu for productive CD8 T cell anti-tumor responses, as collagen-synthesizing macrophages consume environmental arginine, synthesize proline, and secrete ornithine that compromises CD8^+^ T cell function. Thus, a stiff and fibrotic TME may impede anti-tumor immunity not only by direct physical exclusion of CD8^+^ T cells, but also via secondary effects of a myeloid mechano-metabolic programming we identified that creates an inhospitable metabolic milieu for CD8^+^ T cells.

## Introduction

Cancer immunotherapy has revolutionized anti-cancer treatment strategies. However, most cancer patients do not yet respond sufficiently to immunotherapy for it to be broadly curative. Poor responsiveness and/or resistance to immune checkpoint blockade (ICB) has been linked to a failure of immune cells, specifically cytotoxic T lymphocytes (CTL), to infiltrate into the tumor microenvironment (TME) of solid tumors ^1,2^. Recently, a pan-cancer transcriptomic analysis across 20 different cancer types identified a distinct TME subtype characterized by enhanced fibrosis and poor immune infiltration that correlates with poor prognosis and a failure of response to immunotherapy ^3^. Indeed, fibrillar collagen in the TME has been shown to contribute to the physical exclusion of immune cells from the TME, while preventing extracellular matrix (ECM) stiffening improves responses to ICB ^4,5^. However, it remains unclear how collagen and other ECM components regulates anti-tumor immunity and responsiveness to cancer immunotherapy.

Breast cancer progression and aggressiveness are accompanied by increased collagen deposition, macrophage infiltration and TGFβ signaling ^6,7^. TGFβ in the stroma has been shown to drive T cell exclusion from the TME and blunts the response to anti-PD-1/PD-L1 therapies in colorectal and urothelial cancers ^8,9^. This has led to many studies focusing on the role of fibroblasts in TGFβ-mediated ECM-remodeling ^10^, however, fibroblast-targeting therapies, such Sonic Hedgehog (SHH) inhibition have not shown the anticipated anti-tumor efficacy and even worsened tumor progression in patients ^11^. Thus, we hypothesize that alternative mechanisms must be at play contributing to the fibrosis-associated T cell dysfunction in tumors.

It is now well established that metabolism regulates T cell functions and represent a main driver of T cell dysfunction in tumors ^12^. Tumors are metabolically demanding environments in which CTLs compete for critical nutrients ^13^. T cells require glucose and amino acids such as arginine, serine, and glycine for their expansion and anti-tumor activity ^14–17^. Historically, the primary metabolic competition for CTLs in the TME has been thought to be cancer cells, however it has been revealed that tumor infiltrating myeloid cells are the greatest competition for glucose in the TME ^18^. Beyond glucose utilization, tumor infiltrating myeloid cells consume arginine ^19^ via facilitative (SLC7A1/SLC7A2) or anti-port (SLC7A6/SLC7A7) import to then be metabolized by arginase, inducible nitric oxide synthase, or laccase domain-containingL1 ^20,21^. Interestingly, the pan-cancer transcriptomic analysis showed a relationship between fibrosis and abundance of myeloid derived cells that metabolize arginine ^3^, we postulated that changes in myeloid metabolism associated with fibrosis may be an unappreciated determinant of ICB efficacy.

Myeloid metabolism that depletes environmental arginine associates with impaired TCR signaling, proliferation of T cells, production of pro-inflammatory mediators, anti-tumor immunity, and the efficacy of ICB ^17,20,22^. Exogenous supplementation with arginine can improve ICB efficacy, T cell proliferation, and T cell survival ^14,15,17^. However, because T cells import arginine through multifunctional transporters which competitively import a number of structurally similar molecules (e.g., lysine, ornithine, and arginine), the abundance of these competing molecules can dictate whether arginine supplementation improves T cell survival ^14^. Tumor infiltrating myeloid cells also acquire an amino acid metabolism signature marked by SLC25A15 ^23^, the mitochondrial ornithine translocase, which acts downstream of ARG1 for a yet to be determined purpose in this context. Since tumor infiltrating myeloid cells may outcompete T cells for environmental arginine requisite for anti-tumor immunity, it remains to be determined what purpose this metabolic program serves and why it associates with fibrosis. Here, we provide evidences that mechanistically links these independent observations. We demonstrate that breast cancer-associated fibrosis drives a collagen ECM-synthesis program in tumor-associated macrophages that is tightly regulated by TGFβ. We show that this metabolic reprogramming of macrophages is associated with enhanced utilization of arginine and secretion of ornithine into the TME, creating an inhospitable milieu for anti-tumor T cells and ICB.

### Tumor fibrosis and ECM stiffening promote a collagen-ECM synthetic phenotype in myeloid cells

To explore the relationship between fibrosis, stromal stiffening, and myeloid cell phenotypes in the TME we used the FVB MMTV-PyMT (Mouse Mammary Tumor Virus - Polyoma Virus Middle T Antigen) model of mammary tumorigenesis. MMTV-PyMT mice develop mammary tumors by 6-8 weeks that is accompanied by the development of a progressively fibrotic stroma that significantly stiffens as the mice age from 8 to 11 weeks, as confirmed by polarized imaging of picrosirius red stained tissue (Fig.1a; quantified in b, histopathologist scoring Fig. s1a-b) ^7,24^. Our previous work ^7^ revealed that depletion of tumor associated macrophages (TAMs) is sufficient to prevent fibrosis and stromal stiffening observed in 11 week PyMT tumors (Fig. s1c). To explore this finding further, we used flow cytometric analysis to determine if there were changes in the abundance of TAMs between the 8 week and 11 week PyMT tumors (Fig. 1c). Since there were no differences in the total amount of TAMs, we carried out flow-based phenotypic analysis that indicated that the ratio of TAMs expressing higher or lower levels CD11b/CD11c changed significantly in association with the increase in stromal fibrosis (Fig. 1d). Bulk RNAseq comparing the transcriptional profile of TAMs in the nonfibrotic versus the fibrotic tumors (Fig. 1e) revealed a significant increase in the expression of genes associated with interleukin 4 (IL-4) signaling (e.g., *Arg1*, *Vegfa*, and *Mrc1*) (Fig. s1d) and enrichment of both inflammatory and collagen-ECM synthesis and remodeling pathways (Fig. 1f). The findings are consistent with prior *in vitro* studies correlating ECM stiffness with altered TAM phenotype and extend the work to *in vivo* tumors ^6,25^.

**Figure 1:**
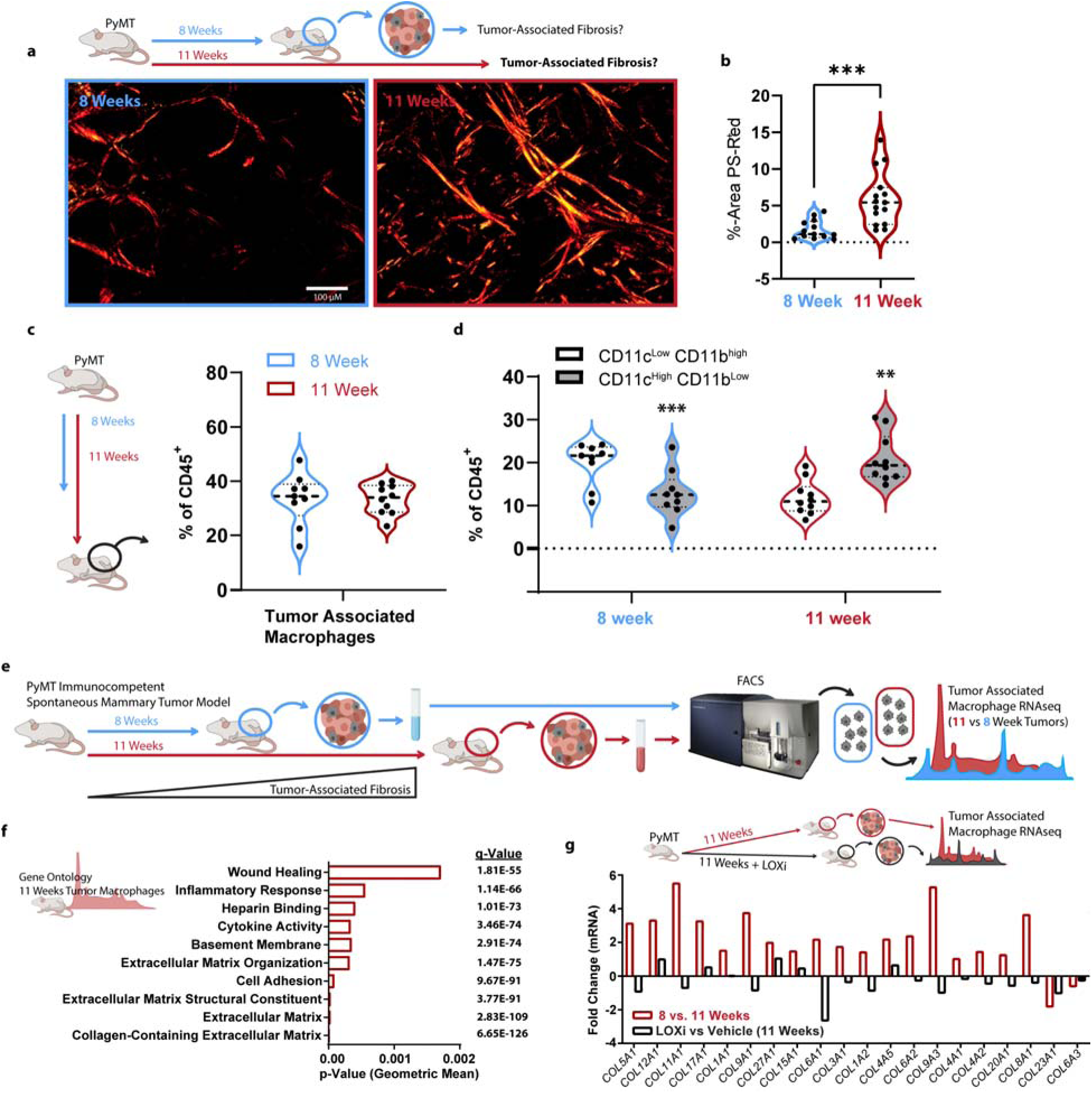
a. Representative birefringence of collagen fibers via polarized light microscopy of picrosirius red (PS-red) stained 8 and 11 week old PyMT mammary tumors, (scale bar: 100 µm). b. % area of PyMT tumor determined to be fibrillar collagen with picrosirius red (PS-red) staining (n=13 or 15). c. Quantification of TAMs found within PyMT mammary tumors from 8 or 11-week old C57BL6/J mice, FACS gating strategy (Supplementary Info: FLOW and RNAseq Gating), (n=9). d. Quantitation of the percentage of CD11c^low^/CD11b^high^ and CD11c^high^/CD11b^low^ of tumor infiltrating CD45^+^ cell types, PyMT mammary tumors from 8 or 11 week old C57BL6/J mice, (n=9). e. Graphical representation of experimental setup for (f). f. Gene Ontology (GO) enriched in TAMs derived from 11 week old PyMT mammary tumors relative to TAMs derived from 8 week old PyMT mammary tumors, (n=5). g. Relative gene expression comparing the top GO category identified in d for TAMs derived from 11 week old PyMT mammary tumors treated with or without LOX-inhibition (LOXi) in the drinking water (∼3 mg/kg/day), (n=5). *Data shown represent ± SEM. *P < 0.05, **P < 0.01, or ***P < 0.005 via two-tailed unpaired Student t test (b-d)*.

To causally link tissue fibrosis and stromal stiffness to the altered TAM phenotype we inhibited lysyl oxidase (LOX) activity with β-aminopropionitrile (LOXi, BAPN) in a cohort of MMTV-PyMT mice to prevent collagen crosslinking and reduce tissue fibrosis and stromal stiffening ^5,24^. We then used RNAseq to transcriptionally profile TAMs isolated from the LOX-inhibited and non-inhibited mice ^24,26^. Bioinformatic analysis revealed that the TAMs isolated from the mice in which stromal stiffening and fibrosis were prevented had reduced levels of collagen-ECM genes (Fig. 1g). Immunofluorescent staining confirmed that inhibiting fibrosis and stromal stiffening reduced the expression of several collagen proteins at the tumor stromal border, including the beaded filament collagen VI (Fig. s1e), previously shown to be deposited in the TME by TAMs ^27–29^. Consistently, TAMs were found in the collagen VI rich regions of the TME (Fig. s1f). We validated this *in vivo* finding by generating mammary fat pad derived fibroblast ECM with and without bone marrow derived macrophages (BMDMs), and observed collagen VI deposition only when BMDMs were present *in vitro* (Fig. s1g).The results suggest that a stiff, fibrotic tumor stroma may trigger a collagen-ECM synthetic myeloid phenotype.

To clarify the clinical relevance of the fibrosis-associated collagen-ECM expression signature by tumor-infiltrating myeloid cells we correlated the expression profiles of myeloid cells isolated from 364 human tumors across 12 cancer types with previously identified tumor-immune states or “archetypes” spanning tissue of origin (Fig. 2a) ^30^. Our analysis revealed that the expression of collagen transcripts in tumor-infiltrating myeloid cells was lowest in Immune Rich and highest in Immune-Stromal and Immune Desert tumor-immune archetypes. The data suggest that expression of a collagen-ECM signature in myeloid cells may be inversely correlated with productive anti-tumor immune responses and patient survival across many tumor types ^30^. Using transcriptomes of 2506 human breast tumor samples (TCGA repository), we next verified that the collagen-ECM signature correlated with IL-4 signaling (Fig. 2b), similar to the association we observed in murine mammary tumor progression and the twelve human tumor types used to define the tumor immune archetypes. Expression of this collagen signature in Stage II and Stage IIIA untreated tumor biopsies also correlated with reduced survival in breast cancer patients (p=0.0346) (Fig. s2). The findings imply myeloid-specific expression of collagen-ECM proteins associates with compromised anti-tumor immunity and poor patient outcome.

**Figure 2:**
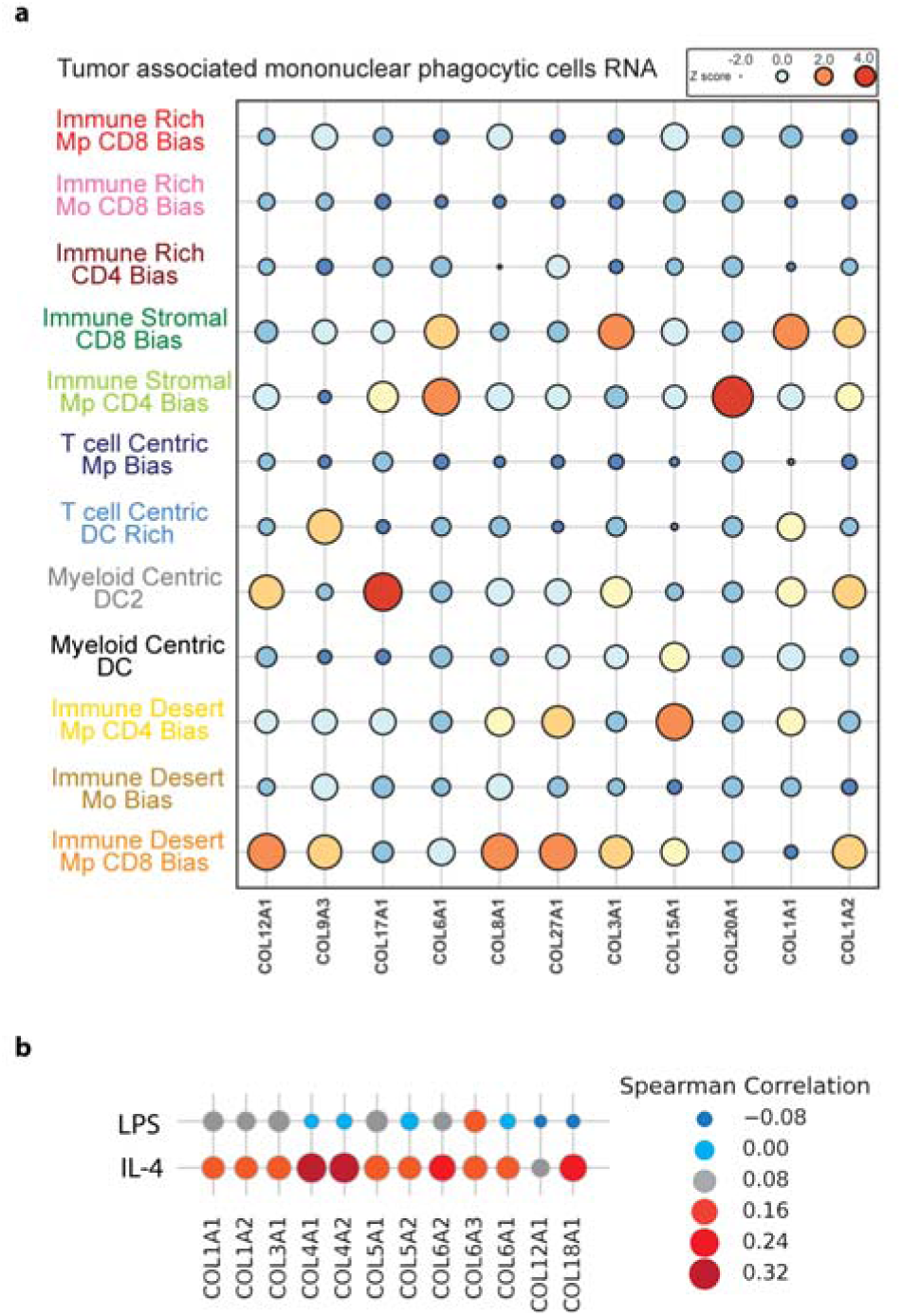
a. Bubble plot of collagen gene expression of the tumor associated myeloid compartment from the Immunoprofiler cohort (IPI) containing 12 different solid tumor types, grouped by cluster/archetype identified in the IPI cohort ^36^. b. Spearman rank correlation of the gene set variation analysis of the top GO category identified in d and macrophage polarization signatures of tumor transcriptomes collected from the TCGA repository (TCGA projects: pancreatic adenocarcinoma, lung adenocarcinoma, glioblastoma multiforme, breast invasive carcinoma, kidney renal clear cell carcinoma).

### A stiff stroma promotes a collagen-ECM synthetic TAM phenotype through altered TGFβ-signaling

We next used an unbiased approach to define what factors control the stiffness-induced, collagen-ECM synthetic TAM phenotype. Promoter motif analysis of the genes comprising the top GO category predicted that they were primarily regulated by SMADs and SP1, a transcriptional response known to occur downstream of TGFβ-signaling (Fig. 3a). We showed previously that CD63 positive myeloid cells secrete TGFβ1 to promote fibroblast-dependent tumor fibrosis ^7^. Autocrine TGFβ-signaling also facilitates the transcriptional identity of tissue resident myeloid populations ^31^. These findings indicate that TGFβ-signaling could promote the stiffness-induced collagen-ECM synthetic myeloid phenotype. Consistently, treating BMDMs with IL-4 and TGFβ1 express collagen VI when the cells were plated on stiff but not soft fibrillar type I collagen-conjugated polyacrylamide gels (Fig. 3b). We also observed a synergistic and independent enhancement of the pro tumor myeloid marker arginase (*ARG1*) and a simultaneous suppression of the wound healing restin-like molecule α (*RELM*α*/RETNLA/FIZZ1*) when the BMDMs were treated with IL-4 and TGFβ1 on the stiff collagen-conjugated polyacrylamide gels (Fig. s3a-b). The data are consistent with our observation in the TAMs isolated from the 11-week PyMT mammary tumors within the stiffened stroma *in vivo* (Fig. s1c). Moreover, we found that the magnitude of TGFβ-induced SMAD signaling depended upon the stiffness of the fibrillar type I collagen the cells interacted with (Fig. s3c). Interestingly, the BMDMs cultured on the stiff fibrillar collagen also produce higher levels of active TGFβ1, determined via TGFβ1-signaling reporter cells (Fig. s3d), which could act in an autocrine manner to reinforce and exacerbate the predominant collagen-ECM synthetic TAM phenotype we observed in the 11-week fibrotic PyMT mammary tumors. The results directly strongly support a positive feedback loop of through TGFβ-signaling in the stiffness-induced collagen-ECM synthetic TAM phenotype.

**Figure 3:**
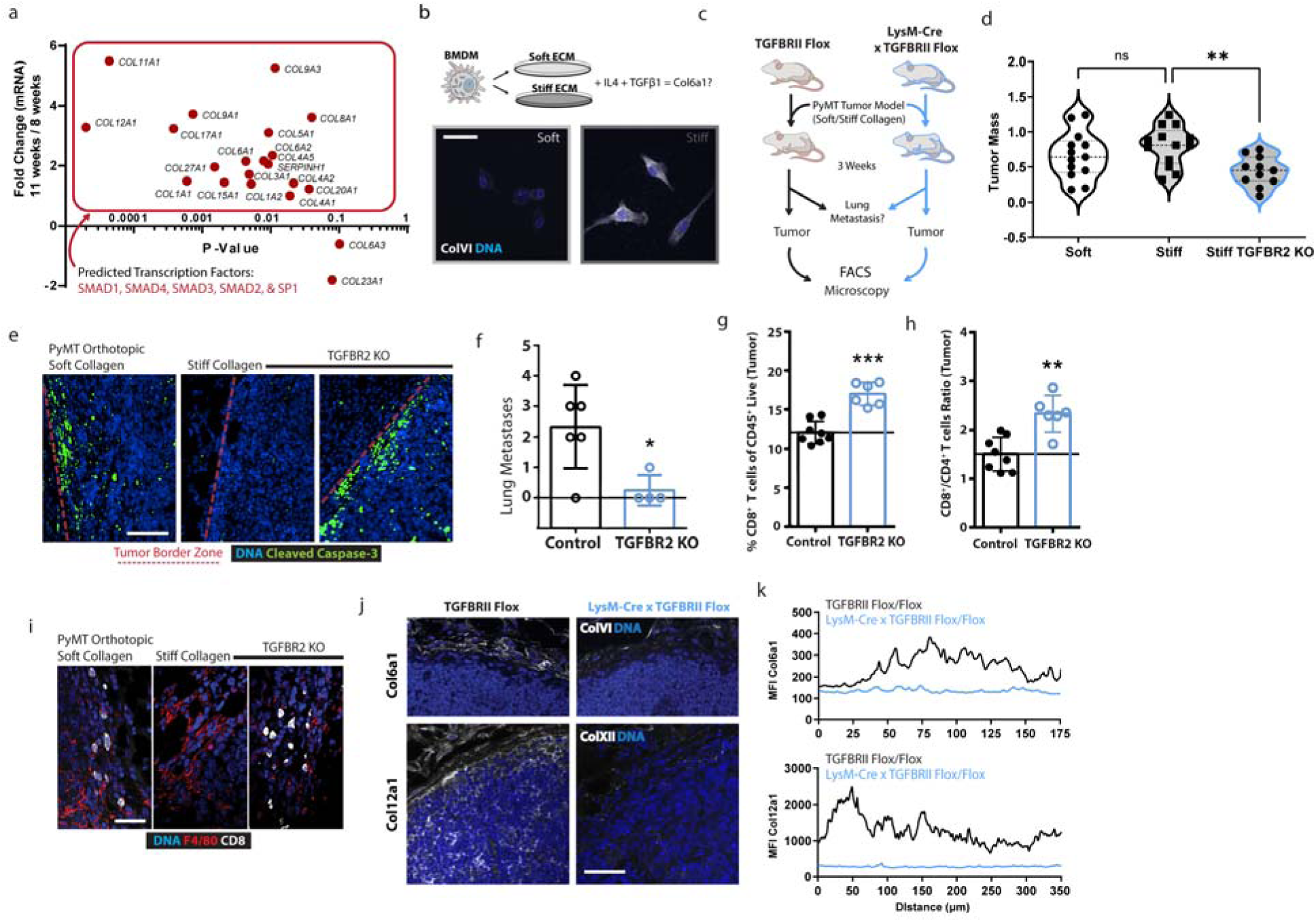
a. XY-plot of relative genes comprising the GO category (collagen-containing ECM) in TAMs derived from 11 week old PyMT mammary tumors relative to TAMs derived from 8 week old PyMT mammary tumors, TRRUST transcription factor enrichment analysis: p-value SAMD1 10^-6^ and SP1 10^-2^.^8^, combined Scores: SMAD1 (127.699827), SMAD4 (90.48497202), SMAD3 (48.37819162), SMAD2 (46.66904886), and SP1 (6.551517203). b. Representative immunofluorescence microscopy of Collagen VI (white) and DNA (blue) of IL4-polarized BMDMs cultured on soft (400 Pa) or stiff (60k Pa) collagen I-coated polyacrylamide hydrogel surfaces treated with 1 ng/mL TGFβ1 for 24 h, (scale bar: 20 µm). c. Graphical representation of experimental setup for (d,e, and g-i). d. Quantification of tumor masses accumulated after 3 weeks of in vivo growth of soft or stiff collagen orthotopic PyMT mammary tumors in TGFβRII^MyeKO^ or control animals, (n=13 or 11). e. Representative immunofluorescence microscopy of cleaved-caspase 3 (red), and DNA (blue) of soft or stiff collagen orthotopic PyMT mammary after 3 weeks of growth in TGFβRII^MyeKO^ or control animals (Scale Bar: 100 µm). f. Quantification of lung metastasis via immunofluorescence microscopy of PyMT staining in lungs of 11 week old spontaneous PyMT mammary tumors TGFβRII^MyeKO^ or control animals (FVB/NJ), (n = 6 or 4). g. Quantification of tumor infiltrating CD8^+^ cells as a percentage of CD45^+^ cells found in stiff collagen orthotopic PyMT mammary after 3 weeks of growth in TGFβRII^MyeKO^ or control animals, FACS gating strategy (Fig. s.xyz), (n=8 or 6). h. Ratio of tumor infiltrating CD8^+^ to CD4^+^ cell types in stiff collagen orthotopic PyMT mammary after 3 weeks of growth in TGFβRII^MyeKO^ or control animals, FACS gating strategy (Fig. s.xyz), (n=8 or 6). i. Representative immunofluorescence microscopy of Collagen VI (white) and DNA (blue) of stiff collagen orthotopic PyMT mammary tumor after 3 weeks of growth in TGFβRII^MyeKO^ or control animals (Scale Bar: 100 µm). i. Representative immunofluorescence microscopy of CD8^+^ (white), F4/80 (red), and DNA (blue) of soft or stiff collagen orthotopic PyMT mammary after 3 weeks of growth in TGFβRII^MyeKO^ or control animals (Scale Bar: 40 µm). j. Quantitation of immunofluorescence microscopy of Collagen VI or XII (white) and DNA (blue) of stiff collagen orthotopic PyMT mammary after 3 weeks of growth in TGFβRII^MyeKO^ or control animals, trace represents the MFI of Collagen VI or XII across 4 transverse sections of the tumor-stromal border (starting ∼50 µm into the stroma and 125-300 µm into the tumor), (n=5 animals averaged into each trace). *Data shown represent ± SEM. **P < 0.01 or ***P < 0.005 via two-tailed unpaired Student t test (f-h) or ANOVA with Tukey test for multiple comparisons (d)*.

To explore the functional links between ECM stiffness and TGFβ-dependent induction of the collagen-ECM synthetic TAM phenotype, we generated a genetically engineered mouse model in which we abrogated TGFβ-signaling in the myeloid cells *in vivo* (LysM-cre x TGFβRII^flox/flox^, TGFβRII^MyeKO^). We then injected PyMT tumor cells embedded within soft and ribose cross-linked and stiffened collagen gels into the fat pads of TGFβRII^MyeKO^ syngeneic host mice, and 3 weeks following injection assayed tumor and stromal ECM phenotype and immune infiltrate (Fig. 3c). Tumor mass was reduced (Fig. 3d) and cell death was increased in the tumor epithelium-stromal border, as indicated by increased levels of cleaved caspase 3 (Fig. 3e) when TGFβ-signaling was abrogated in the tumor associated myeloid cells that developed in the stiff collagen matrices. Next, to explore the impact of myeloid cell TGFβ-signaling more rigorously on mammary tumor behavior we next used a spontaneous tumor model in which we crossed FVB PyMT mice with LysM-Cre/TGFβRII floxed mice. While ablating TGFβ-signaling in the myeloid cells had only a modest impact on primary tumor development in the spontaneous PyMT model (data not shown), we quantified a significant reduction in lung metastasis (Fig. 3f). The results underscore the physiological impact of normalizing myeloid cell function through ablating TGFβ-signaling on primary breast tumor phenotype and metastasis.

TAMs can repress the function and viability of tumor-associated CD8^+^ T cells modifying tumor phenotype and influencing metastasis ^32,33^. Nevertheless, immunostaining indicated that myeloid cell infiltration was similar between the PyMT orthotopic tumor models in C57BL6/J mice (data not shown). We did, however, quantify an increase in the total number of live CD8^+^ T cells and found an enhanced ratio of CD8^+^/CD4^+^ T cells within the orthotopic tumors that developed in the stiff tumors when TGFβ−signaling was ablated in TAMs (Fig. 3g-h). Consistently, using immunofluorescence we observed more CD8^+^ T cells infiltrating into stiff collagen TGFβRII^MyeKO^ syngeneic PyMT tumors (Fig. 3i). Furthermore, we observed a dramatic reduction in the level of collagen VI at the tumor-myeloid-stromal boundary and collagen XII at the boundary and throughout the TME when TGFβ-signaling was impeded in the myeloid cells in the stiff tumors (Fig. 3j, quantified in k). The data imply that abrogating TGFβ-signaling in TAMs prevents the stiffness-dependent induction of their collagen-ECM synthetic phenotype. Importantly, normalizing the TGFβ-dependent TAM phenotype also repressed tumor aggression and metastasis and restored CD8^+^ T cell infiltration into the tumor, despite the presence of a stiff stroma. The findings suggest a stiff, fibrotic tumor stroma likely compromises CD8^+^ T cell infiltration into the tissue by fostering a collagen-ECM synthetic phenotype in the TAM population.

### Mechanosignaling and TGFβ control collagen-synthetic metabolic programming in macrophages

Thus far, we have shown that tumor progression promotes a TGFβ-driven feed-forward fibrotic program in TAMs that is correlated with lower CD8^+^ T cell infiltration and dysfunctional anti-tumor immunity. The next question we sought to answer is how does fibrosis restrict T cell mediated anti-tumor immunity, is it the product of a physical barrier or a byproduct of the metabolic consequences of fibrosis? Activated T lymphocytes have unique metabolic demands to support their effector functions and survive in the TME ^14^. Specific metabolic programs are essential for immune cells to achieve discrete phenotypes or functions ^34–36^, and the available metabolites can dictate whether a cell can utilize a particular metabolic pathway to achieve an optimal phenotype or function ^14,37–39^. Since ECM stiffness exerts significant effects on cellular metabolism ^40^, we profiled polar metabolites in spontaneous PyMT tumors in which stromal stiffness and fibrosis were prevented through inhibition of LOX crosslinking. Our objective was to identify whether the metabolic milieu was regulated by tissue fibrosis and stromal stiffness and to determine whether this could influence the phenotype or function of the infiltrating immune cells. We found that preventing collagen crosslinking and stromal stiffening via LOX-inhibition was accompanied by a ∼4-fold increase in the concentration of arginine in the TME (Fig. 4a). These data are consistent with previous work showing the interstitial fluid of tumors is depleted of arginine relative to normal tissues ^41^.

**Figure 4:**
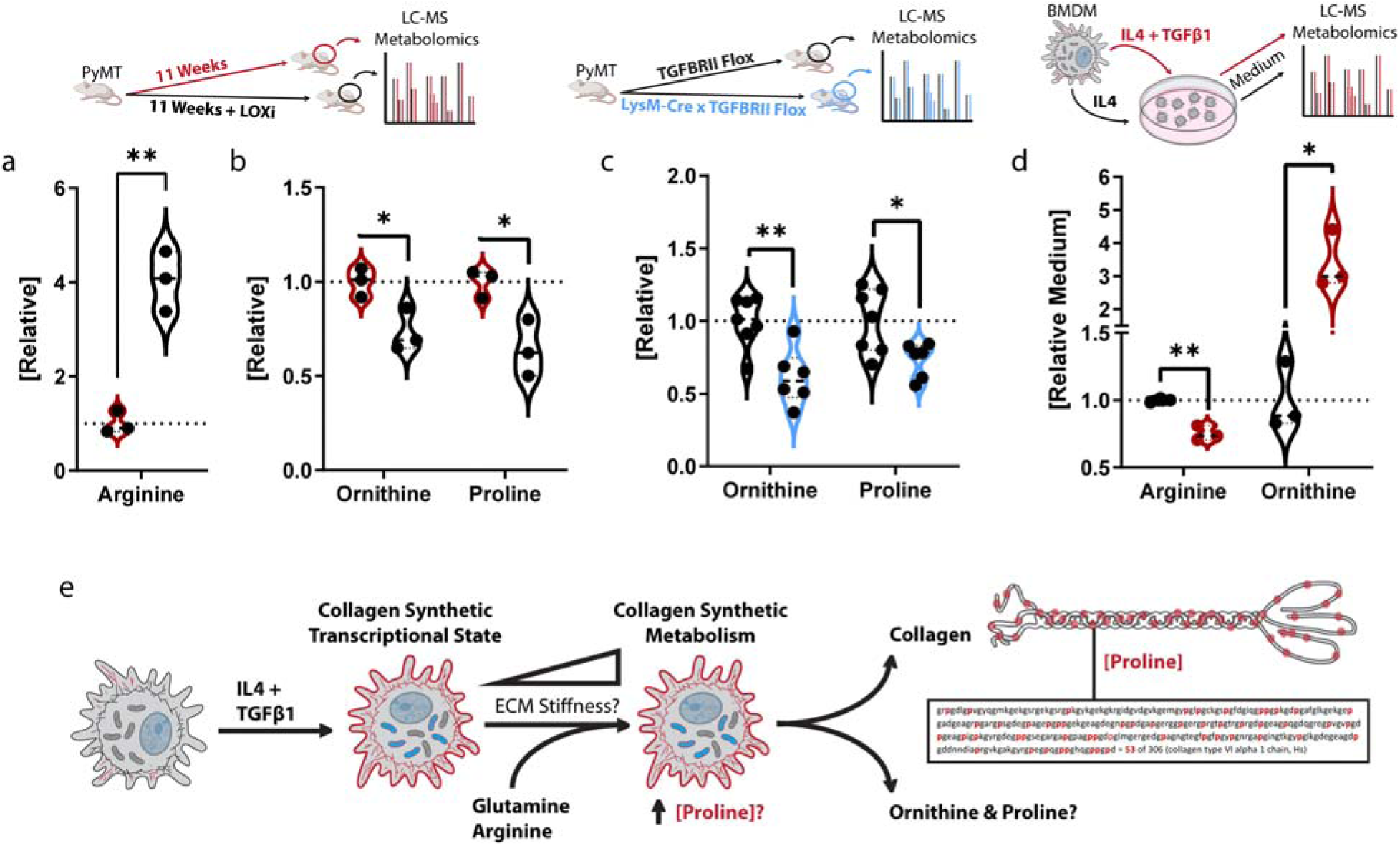
a. Relative arginine concentration of 11 week old PyMT mammary tumors treated with or without LOX-inhibition (LOXi) in the drinking water (∼3 mg/kg/day), LC-MS analysis, (n=3). b. Relative ornithine and proline concentrations of 11 week old PyMT mammary tumors treated with or without LOX-inhibition (LOXi) in the drinking water (∼3 mg/kg/day, LC-MS analysis, (n=3). c. Relative ornithine and proline concentrations of stiff collagen orthotopic PyMT mammary tumors in TGFβRII^MyeKO^ or control animals for 3 weeks, LC-MS analysis, (n=7 or 6). d. Relative medium concentrations of arginine and ornithine from culture with IL4-polarized BMDMs treated with or without 1 ng/mL TGFβ1 for 22 h, swapped for fresh medium for 2 h which was measured via LC-MS analysis, (n=3). e. Graphical depiction of the observed relationship between TGFβ1 signaling and ECM stiffness in the context of collagen synthesis and proline metabolism. *Data shown represent ± SEM. *P < 0.05 and **P < 0.01 via two-tailed unpaired Student t test (a-d)*

Myeloid cells expressing high levels of *ARG1* have been implicated as major consumers of arginine in the TME ^42,43^. Collagen contains a disproportionally high abundance of proline (∼8 fold higher than proteome median), which can be synthesized from arginine or glutamine. Tumor metabolomics revealed that corresponding to the increased levels of arginine there was a commensurate decrease in two products of arginine metabolism, proline and ornithine, in the PyMT mice in which collagen crosslinking and stromal stiffening and tissue fibrosis were prevented (Fig. 4b). The findings provide one explanation for why supplementation with arginine enhances collagen deposition and tensile properties of healing wounds ^44,45^ and has previously been shown to enhance the clinical efficacy of treating “hard-to-heal” wounds in elderly and diabetic patients ^46^. TGFβ-signaling enhances proline synthesis ^47^ via glutamine and arginine metabolism ^48^. Because there is evidence that there may be a direct relationship between arginine depletion and ornithine enrichment in the TME ^41^ we next sought to determine if myeloid TGFβ-signaling affected the proportions of arginine:ornithine:proline in tumors. We assayed polar metabolites in the TME of TGFβRII^MyeKO^ tumors and their corresponding controls. We found that while levels of arginine were unaffected (data not shown), ornithine and proline levels were significantly reduced in TGFβRII^MyeKO^ tumors (Fig. 4c), implicating TAM-TGFβ signaling as a significant regulator of the intratumoral levels of these metabolites. To determine whether TGFβ-signaling can directly influence myeloid utilization of exogenous arginine, we assayed IL-4 stimulated BMDMs interacting with a stiff matrix that were treated with or without TGFβ. Data revealed that TGFβ enhanced arginine uptake, as well as ornithine efflux into the culture medium (Fig. 4d-e).

We then tested if ECM stiffness and TGFβ synergistically enhanced arginine conversion into proline by supplementing IL-4-polarized BMDMs with ^13^C_6_-arginine and measuring proline (m+5) (which indicates the direct flow of carbon derived from arginine into *de novo* proline synthesis) (Fig. 5a-b). Proline synthesis rates from arginine metabolism (Fig. 5b) and steady-state proline-OH levels (Fig. s4a) were both synergistically enhanced by TGFβ and ECM stiffness, whereas intracellular ornithine levels (Fig. 5c) were only elevated by ECM stiffness, and steady-state proline levels were specifically elevated in response to TGFβ (Fig. s4b). Because de novo proline synthesis can also be fed by glutamine metabolism, which is mechano-sensitive ^49^, we assayed if ECM stiffness influenced glutamine conversion into proline. Strikingly, we found that stiff fibrillar collagen caused ∼5-fold higher incorporation of ^13^C into proline from ^13^C_5_-glutamine (Fig. 5d) indicating that the two primary pathways implicated in *de novo* proline synthesis are both mechanosensitive and TGFβ-sensitive. To determine whether stiffness influenced myeloid utilization of exogenous arginine or glutamine, we assayed IL-4 stimulated BMDMs interacting with a soft or stiff matrix treated with or without TGFβ and quantified the metabolite composition of the medium. We observed that the primary metabolite consumed from the medium was arginine, and the primary metabolites that accumulated in the medium were proline and ornithine (Fig. 5e).

**Figure 5:**
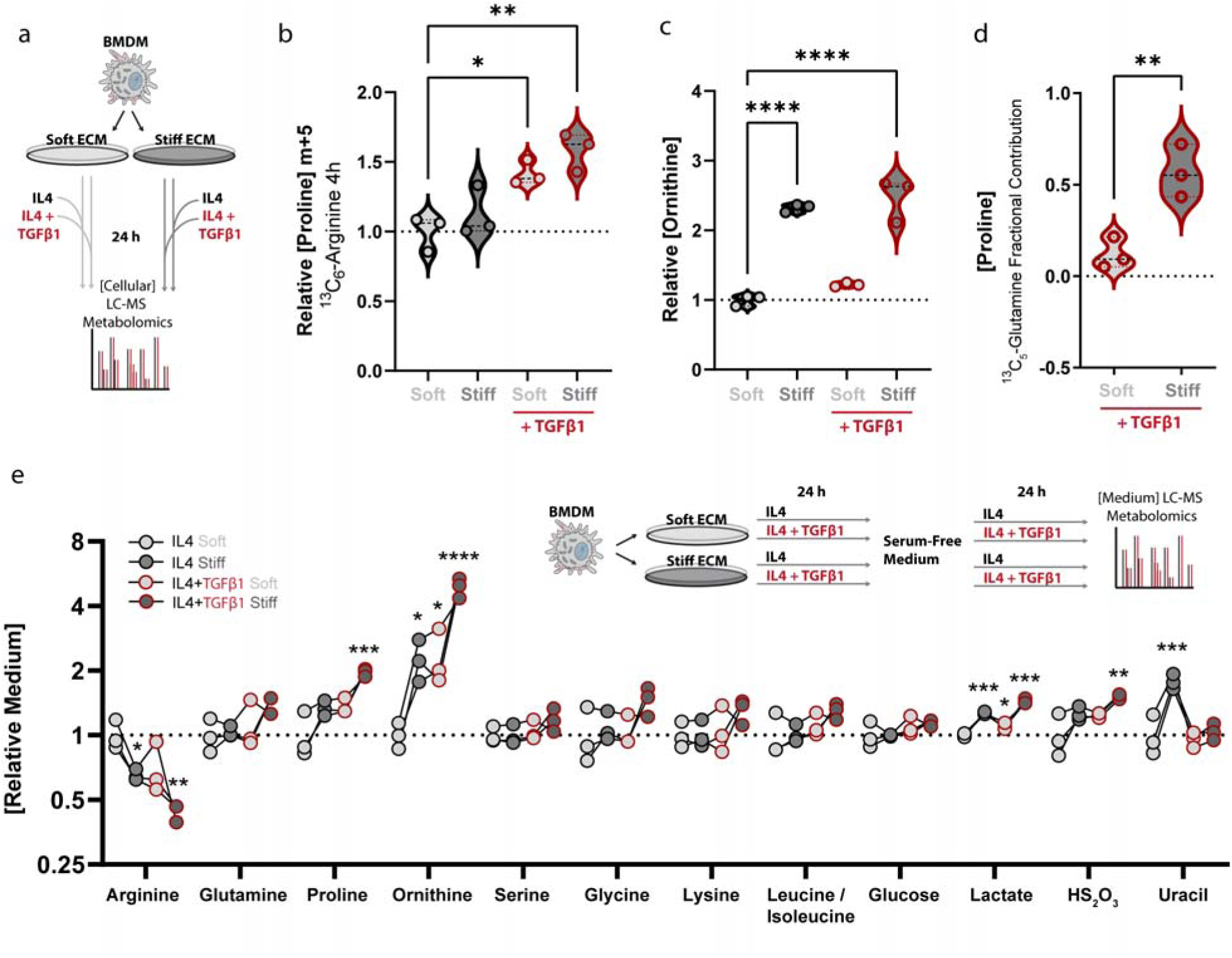
a. Graphical representation of experimental setup for (f-g). b. ^13^C tracing of ^13^C_6_-arginine metabolism into ^13^C_5_-proline in BMDMs cultured on soft (400 Pa) or stiff (60k Pa) collagen I-coated polyacrylamide hydrogel surfaces in medium containing ^12^C_6_-arginine, treated with or without 1 ng/mL TGFβ1 for 20 h, swapped for fresh medium containing ^13^C_6_-arginine for 4 h, BMDMs were harvested and measured via LC-MS, (n=3). c. Relative ornithine concentration in BMDMs cultured on soft (400 Pa) or stiff (60k Pa) collagen I-coated polyacrylamide hydrogel surfaces with or without 1 ng/mL TGFβ1 for 24 h, measured via LC-MS, (n=3). d. Fractional contributions (i.e., all isotopologues) of ^13^C derived from ^13^C_5_-glutamine in BMDMs cultured on soft (400 Pa) or stiff (60k Pa) collagen I-coated polyacrylamide hydrogel surfaces in medium containing ^12^C_5_-glutamine, treated with or without 1 ng/mL TGFβ1 for 22 h, swapped for fresh medium containing ^13^C_5_-glutamine for 2 h, BMDMs were harvested and measured via LC-MS, (n=3). e. Relative concentrations of metabolites in culture medium of IL4-polarized BMDMs cultured on soft (400 Pa) or stiff (60k Pa) collagen I-coated polyacrylamide hydrogel surfaces treated with or without 1 ng/mL TGFβ1 for 24 h then swapped for fresh serum-free medium for 24 h which was measured via LC-MS analysis (n=3). *Data shown represent ± SEM. *P < 0.05, **P < 0.01, or ****P < 0.0001 via one-way ANOVA with Tukey test for multiple comparisons (b-e)*.

It has been proposed that proline synthesis serves as vent for excess mitochondrial reducing equivalents (e.g., NADH and NADPH) when mitochondrial respiration is suppressed. Mitochondrial oxidative flux and the activity of pyrroline-5-carboxylate reductase 1 (PYCR1), required to synthesize proline in the mitochondria are mechanosensitive ^40,50^. We found that TGFβ suppressed mitochondrial respiration (Fig. s4c) and NADH levels were synergistically enhanced by TGFβ and ECM stiffness (Fig. s4d). Although we did not observe an ECM stiffness dependent enrichment of PYCR1 in the mitochondria of BMDMs, we did observe an overall increase in mitochondrial content in the BMDMs interacting with the stiff fibrillar collagen ECM. The findings imply that the increased proline synthesis we detected in IL-4-polarized BMDMs cultured on a stiff ECM was primarily driven by a compensatory response to elevated levels of NADH, similar to what has been found in fibroblasts ^47^.

### Ornithine impairs CD8 CTL metabolism and anti-tumor activity

Commensurate with arginine consumption in TAMs and IL-4-polarized BMDMs, we quantified an increase in environmental ornithine in the TME of the PyMT tumors, as well as in the culture medium of TGFβ-stimulated IL-4-polarized BMDMs interacting with a stiff ECM. Arginine is an essential metabolite for activated lymphocyte survival and anti-tumor responses^14^. In fact, conditioning tumors with arginine synthetic bacteria, that consume ammonia to synthesize arginine, potentiates cancer immunotherapies reliant on activated CTLs ^15^. Models of T cell activation using CD3/CD28 antibodies have demonstrated that activation triggers distinct metabolic programing ^14,38,51^. To explore how ornithine affects CD3/CD28-activated CD8^+^ CTL metabolism, we titrated the concentration of ornithine relative to arginine (∼1:1, 3:1, and 9:1 that mimics the effect observed in tumors) and measured intracellular polar metabolites with mass-spectrometry based metabolomics. We found that after 24 hours of activation, increased concentrations of ornithine resulted in a decrease in ATP and elevated levels of intracellular ornithine, its downstream metabolites (e.g., citrulline) (Fig. s5a). While it is not clear why arginine is the favored metabolic substrate in activated CTLs, if arginine metabolism is suppressed and ATP levels fall, translation (high ATP demand) could be impeded, and the synthesis of effector proteins stalled retarding anti-tumor programming. Peak utilization of arginine has been shown to occur ∼ 72h after activation. Accordingly, we assayed the metabolome of CD8^+^ CTLs after 3 days of culture in an environment containing a molar ratio of ∼3:1 ornithine:arginine (Fig. 6a). Results revealed that nearly all metabolite pools were suppressed by environmental ornithine, with the exception of oxidized glutathione (GSSG) which negatively affects survival and inflammatory programming of CTLs ^52,53^.

**Figure 6:**
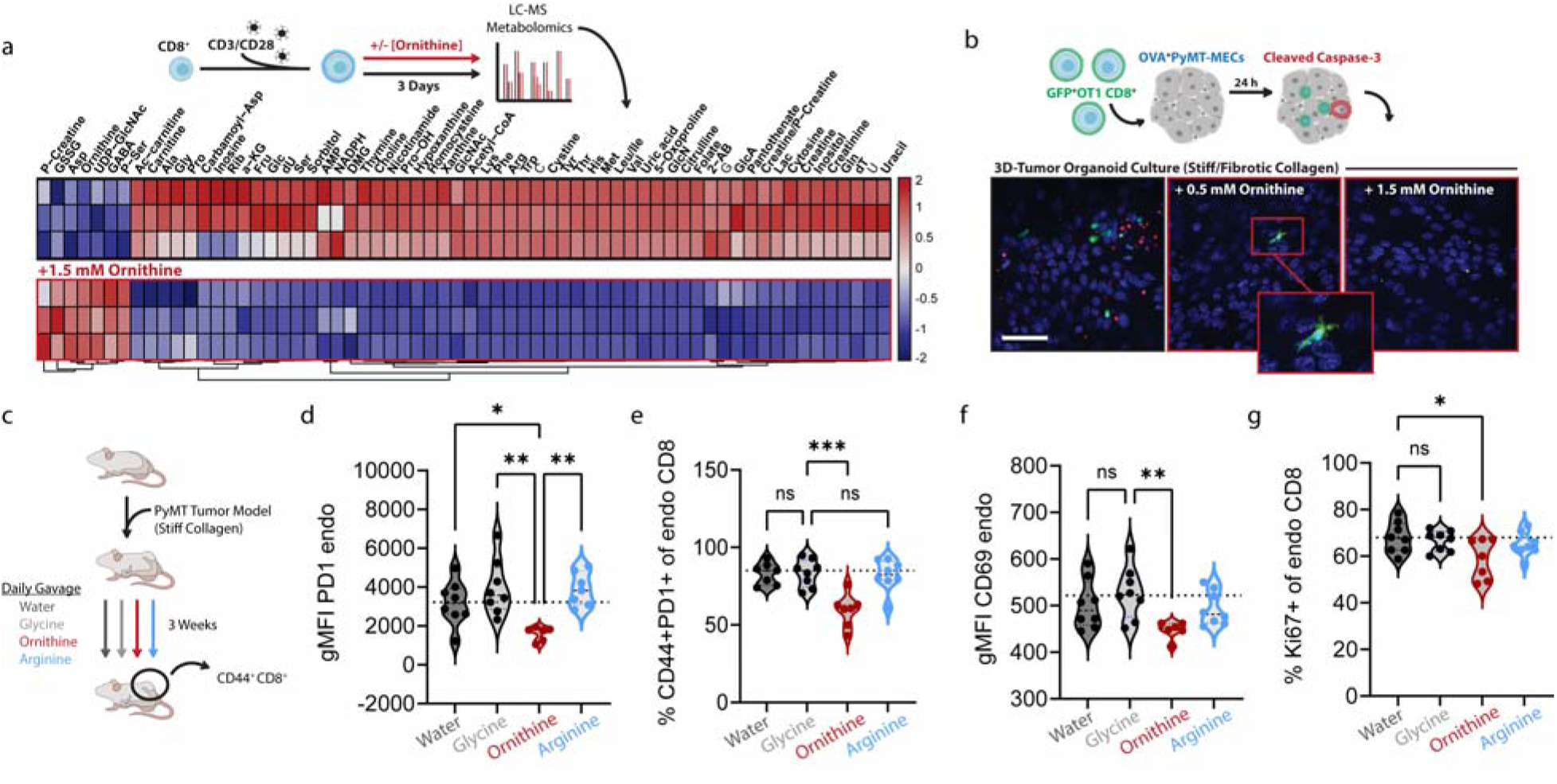
a. Heat map of relative metabolite levels of CD3/CD28-activated CD8^+^ CTLs cultured for 72 h in medium containing a molar ratio of 3:1 ornithine:arginine, medium refreshed every 24 h, LC-MS analysis. (n=3 biological replicates) b. Representative immunofluorescence microscopy of stiff-collagen OVA-PyMT tumor organoids challenged with GFP^+^ OT-I CTLs (green) for 24 h in medium containing a molar ratio of 1:1 [0.5mM] or 3:1 [1.5 mM] ornithine:arginine, cleaved-caspase 3 (red) and DNA (blue), (Scale Bar: 40 µm). c. Graphical description of the experimental setup for d-f. d. Mean fluorescent intensity of PD1 staining of endogenous CD8^+^ CTLs isolated from stiff collagen PyMT tumors after 3 weeks of growth in C57BL6/J mice gavaged daily with 2 g/kg glycine, ornithine, arginine, or water (100 µL), (n=8). e. Percentage of PD1^+^ endogenous CD8^+^ CTLs isolated from stiff collagen PyMT tumors after 3 weeks of growth in C57BL6/J mice gavaged daily with 2 g/kg glycine, ornithine, arginine, or water (100 µL), (n=8). f. Mean fluorescent intensity of CD69 staining of endogenous CD8^+^ CTLs isolated from stiff collagen PyMT tumors after 3 weeks of growth in C57BL6/J mice gavaged daily with 2 g/kg glycine, ornithine, arginine, or water (100 µL), (n=8). g. Quantification of KI67 incorporation into endogenous CD8^+^ CTLs isolated from stiff collagen PyMT tumors after 3 weeks of growth in C57BL6/J mice gavaged daily with 2 g/kg glycine, ornithine, arginine, or water (100 µL) and stimulated ex vivo with PMA [50 ng/mL] and ionomycin [500 ng/mL], (n=8). *Data shown represent ± SEM. *P < 0.05 or **P < 0.01 via one-way ANOVA with Tukey test for multiple comparisons (d-g)*.

To determine if ornithine levels can affect antigen-specific CD8^+^ CTL anti-tumor responses, we utilized a cell line derived from the PyMT-mCherryOVA model ^54^ that express the model antigen chicken ovalbumin (OVA) and transgenic OT-I CD8^+^ T cells that specifically recognize the OVA-derived SIINFEKL peptide. OVA-PyMT tumor organoids embedded within a ribose-stiffened 3D collagen gel or cultured as a monolayer on top of rigid glass coverslips were challenged with GFP^+^ OT-I cells in the presence of varied concentrations of ornithine (Fig. 6b and s5b). The cultures were then assayed for markers of death induction by staining for cleaved caspase 3. Tumor cells cultured in the presence of ornithine appeared to be resistant to OT-I-mediated death (e.g., cleaved caspase-3^+^) and the only GFP^+^ OT-I cells we could find under either culture condition were already dead and fragmented (∼ 1-3 um fragments with no observable nuclei) or were in the process of dying, as indicated by positive staining for caspase 3.

Because *in vitro* models do not consistently recapitulate living tissues, we next tested if dietary supplementation could be used to raise the concentration of circulating ornithine or arginine in an orthotopic MMTV-PyMT mammary tumor model, and if this influenced tumor phenotype and CD8^+^ CTL function (Fig. s5c). We injected PyMT tumor cells embedded within ribose cross-linked and stiffened collagen gels into the mammary fat pads of syngeneic host mice and conducted daily oral gavages of water, glycine, ornithine, or arginine for three weeks. The glycine supplementation was included as an approximate caloric control for amino acid supplementation. *In vivo* supplementation with ornithine raised the concentration of plasma ornithine ∼12-fold, NAD^+^ 4-fold, but it did not affect citrulline (Fig. s5d). Supplementation with arginine elevated circulating arginine and ornithine ∼3.5-fold and similarly did not significantly affect citrulline levels. As a control, we tested both water and glycine, which did not affect circulating levels of ornithine or arginine, although we did observe a modest increase in citrulline levels when we supplemented with glycine (Fig. s5d). Dietary supplementation with ornithine, arginine, or glycine did not affect orthotopic PyMT primary tumor size (data not shown), but we did quantify more circulating tumor cells in the mice that were supplemented with ornithine (Fig. s5e). Ornithine supplementation reduced the proportion of activated CD44^+^ PD1^+^ endogenous tumor-infiltrating CD8^+^ CTLs as well as the expression levels of PD-1 and CD69, whereas arginine supplementation did not, in spite of also elevating circulating ornithine levels (Fig. 6c-f and s5f). Moreover, *ex vivo* re-stimulated CD8^+^ CTLs isolated from these tumors were less proliferative as indicated by reduced Ki67 staining (Fig. 6g). These findings suggest ornithine may impair CD8^+^ CTL activation in the TME.

PD1 expression on CD8^+^ CTLs typifies ongoing recognition of tumor antigen that may be responsive to checkpoint blockade-based immunotherapies ^55^ and ornithine supplementation strongly decreased PD1 expression in CTL infiltrating PyMT tumors. Accordingly, we sought to test if PyMT tumor responsiveness to PD1 blockade was affected by the elevated circulating levels of ornithine. While ornithine, arginine, or glycine supplementation alone did not affect orthotopic PyMT tumor growth, they did affect tumor growth rates in the mice that were treated with anti-PD1 checkpoint blockade (Fig. 7a-b). Ornithine supplementation however, abolished any beneficial effects of the anti-PD1 blockade such that the tumors grew to their humane endpoint as quickly as isotype-treated control animals (Fig. 7c). By contrast, checkpoint blockade in the context of arginine and glycine supplementation exerted a ∼33% increase in the median survival time of the treated mice. These data suggest that fibrotic tumors, which are enriched in ornithine via collagen-ECM synthetic TAMs, may impede anti-tumor CTL responses by creating an inhospitable metabolic milieu for their anti-tumor functions in the TME (Fig. 7d).

**Figure 7:**
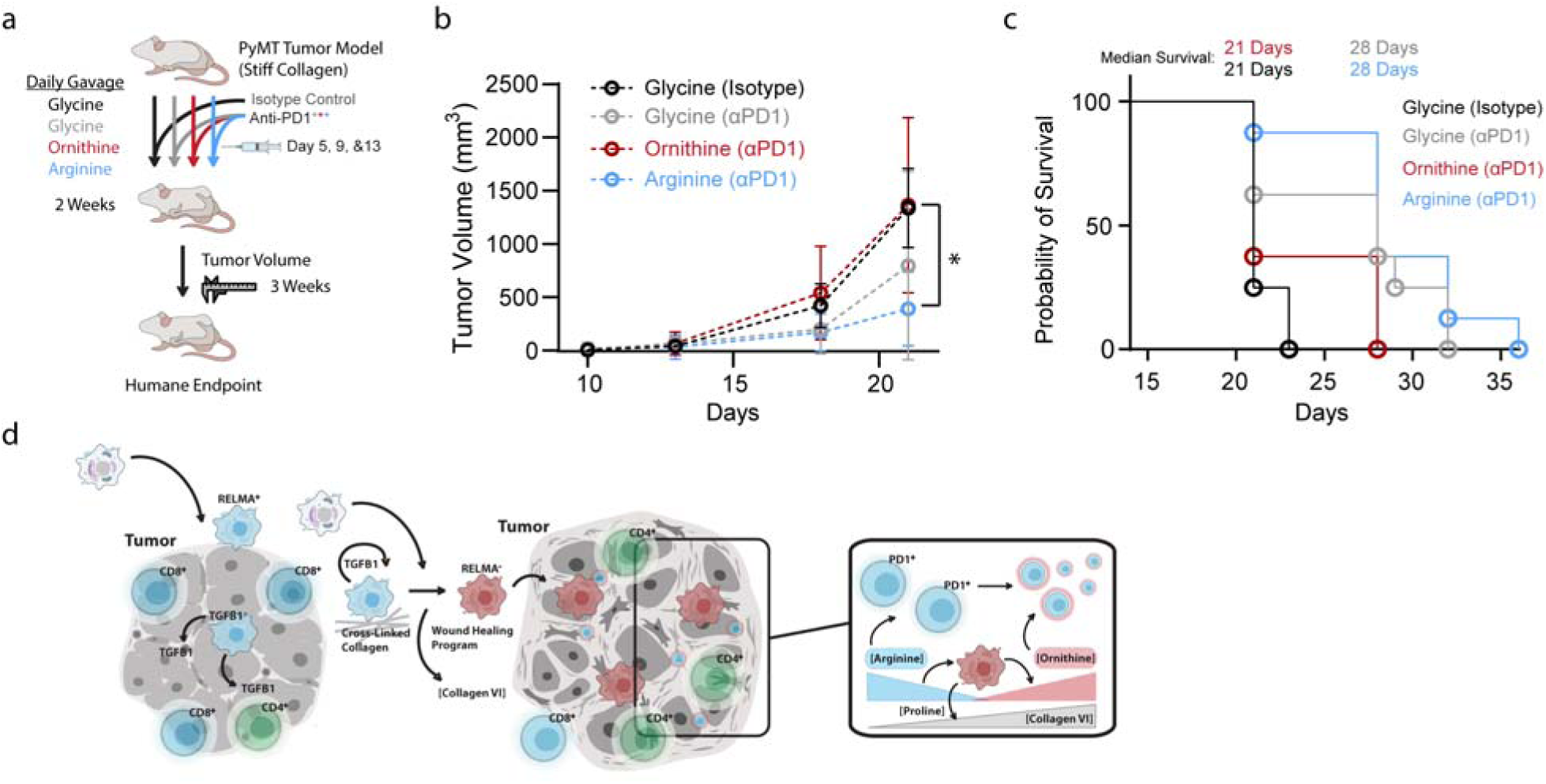
a. Graphical description of the experimental setup for h-i. b. Tumor volume measurements of stiff collagen PyMT tumors over 4 weeks of growth in C57BL6/J mice gavaged daily with 2 g/kg glycine, ornithine, or arginine (100 µL) for the first 14 days and treated with isotype control or anti-PD1 blocking antibodies on day 5, 9, and 13, humane endpoint indicated by horizontal dotted line and white background (n=8). c. Kaplan-Meier survival curves of stage of experiment depicted in h. Mean survival of glycine treated isotype control and ornithine treated anti-PD1: 21 days, Mean survival of glycine and arginine treated anti-PD1: 28 days. d. Graphical representation of the identified relationship between the TME, infiltrating myeloid cells, mechano-metabolic programming, and CTLs in the TME.

## Discussion

We found that the stiff fibrotic tumor stroma synergizes with TGFβ to induce a collagen-ECM synthetic phenotype in TAMs in the TME. Our studies indicate that collagen-ECM synthetic TAMs generate an unfavorable ornithine-rich metabolic barrier, which compromises CD8^+^ T cell function and limits ICB responsiveness. Our results expand that prevailing dogma that cancer fibrosis acts as a physical barrier that surrounds and impedes CD8^+^ T cell migration into the tumor to create an “immunologically cold” TME ^8,9^. Our observations offer an additional explanation for these results, in which the fibrotic stroma rewires the tumor-associated TAM phenotype to create a noxious metabolic microenvironment around the tumor that compromises the function of the CTLs. Indeed, while prior work showed that inhibiting LOX-dependent cross-linking and stromal stiffening in PyMT mice increased T cell motility and improved responses to ICB, using the same model we showed that LOX-inhibition reverts the collagen-ECM synthetic TAM phenotype to normalize the TME metabolic milieu thereby restoring anti-tumor immunity ^4,5^. Thus, a stiff and fibrotic TME may impede anti-tumor immunity not only by direct physical exclusion of CD8^+^ T cells, but also through a secondary effect of a macrophage mechano-metabolic programming we identified that creates an inhospitable metabolic milieu for CD8^+^ T cells.

TAMs can impair CTL anti-tumor responses and diminish the efficacy of checkpoint blockade ^32,56^. To address this, strategies have been developed to ablate TAMs with varying clinical successes, possibly due to the high heterogeneity and plasticity of TAM ^57,58^. Indeed, although distinct TAM populations have previously been characterized with flow cytometry based on expression levels of CD11b/CD11c ^68,69^, scRNAseq efforts reveal that TAMs possess much greater phenotypic diversity ^23,59–61^. Thus, while subsets of TAMs exist that can compromise anti-cancer immunotherapy by releasing CTL-suppressive cytokines, engaging in metabolism that competes for CTL demands, and stimulating fibrosis to impede CTL tumor-infiltration ^7,9,62^, not all TAMs exert repression on the adaptive immune system or promote tumor growth. Recent studies have shown that macrophage heterogeneity and function in the TME may be associated with specific metabolic programs ^23^. Indeed, we identify the stiff, fibrotic stroma of the tumor as a key regulator of the pro-tumor phenotype through TGFβ signaling and expand this characterization to include its collagen-ECM synthetic ornithine secretory arginine uptake behavior. Our data establish the arginine/ornithine metabolism of TAMs as a key determinant of the cytotoxic activity adaptive immune cells, particularly CTLs in the context of ICB ^63^.

Early infiltrating TAMs promote the development of tumor-associated fibrosis by conditioning the TME with TGFβ1, which acts on stromal fibroblasts to synthesize fibrillar collagens and collagen crosslinking enzymes leading to a stiffened TME ^7^. The stiffened, fibrotic ECM in turn recruits TAMs ^6,64^, promotes tumor aggression ^65^, and metabolically reprograms tumor cells ^40^ and infiltrating immune cells. Our results demonstrate that the infiltrating TAMs themselves synthesize and secrete ECM proteins, including collagen VI and XII which can exert pleiotropic effects on the organization and stiffness of the tumor epithelium and noncellular and cellular stroma ^66,67^. Thus, while cancer-associated fibroblasts are the primary source of fibrillar collagen in the TME, TAMs contribute specific collagens to the TME, such as the beaded filament collagen VI which forms microfibrillar networks that entrain epithelial basement membranes to interstitial ECM and facilitate wound healing, scar tissue formation, re-epithelialization, and metastatic progression ^68,69^. The relative contribution of TAM-deposited ECM proteins to tumor fibrosis and its role in creating spatial ECM and cellular heterogeneity clearly merit further investigation.

Therapeutic approaches intended to treat fibrosis in the TME have not yet yielded clinical success, possibly because they were not combined with ICB, targeted inappropriate stromal cell types, or were based upon inaccurate assumptions. For instance, cancer-associated fibrosis is thought to drive tumor aggression in part because it serves as a physical barrier to CLT infiltration ^4,5^. Yet, T cells have been shown to migrate through sub-micron spaces when extravasating into inflamed tissues or through engineered pore sizes much smaller than the highly cross-linked fibrillar collagen matrices observed in tumors, implying the dense collagen-rich ECM in tumors is unlikely to impede T cell infiltration ^70–73^. Our findings suggest that the fibrotic TME distorts antitumor immunity though alterations in myeloid metabolism, so that strategies designed to remediate the metabolic microenvironment associated with fibrosis may be a more effective anti-tumor strategy. Indeed, the increased experimental pancreatic adenocarcinoma (PDAC) aggression induced by ablating αSMA positive fibroblasts and depleting fibrillar collagen was accompanied by a massive infiltration of pro-tumorigenic myeloid cells, which our studies suggest may have driven tumor aggression and compromised anti-tumor immunity through their aberrant metabolism ^74^. Consistently, therapies that reduce myeloid infiltrate in the TME have beneficial effects with and without ICB ^75,76^, whereas treatments intended to target fibroblasts in the TME have thus far shown limited clinical benefit ^11^. As we and others illustrated through metabolic supplementation or the use of Arg-ECN to enrich the TME with arginine offers a viable approach to improve anti-tumor immunity. Additionally, since a SLC25A15 expressing subset of TAMs are likely to be the major contributor to arginine depletion and ornithine enrichment in the TME ^23^, identifying ways to selectively ablate this TAM population may be sufficient to prevent myeloid pro-tumor programming, while simultaneously maintaining anti-tumor myeloid populations in the TME. Overall, these types of approaches might have higher potential to overcome recent failures encountered with anti-stromal therapies due to factors including non-specificity, poor penetration or stromal fibroblast heterogeneity ^11^.

Arginine supplementation can potentiate anti-tumor immunity and the efficacy of checkpoint blockade therapies ^15^. However, ingested arginine can be metabolized in the gut or liver, generating ornithine, which based upon our studies could exert suppressive effects on CTL proliferation and activation ^77^. Furthermore, our results indicate that although arginine supplementation can exert beneficial effects on CTL function it also simultaneously increases the circulating concentration of ornithine which would potentially compromise its beneficial impact on anti-tumor immunity. Consistently, chronic viral infections of the liver increase circulating ornithine levels which impede virus-specific T cell expansion, indicating that ornithine is a systemic immunomodulatory metabolite for CTLs ^78^. Accordingly, therapies that locally increase arginine through consumption of ornithine as opposed to systemic arginine supplementation may offer a greater ability to enhance checkpoint blockade therapies and promote anti-tumor immunity. Towards this objective local synthesis of arginine by a synthetically-engineered strain of Escherichia coli Nissle 1917 (ECN) that was designed to consume environmental ammonia and excrete arginine in the TME was recently shown to improve checkpoint blockade ^15^. Due to the nature of the metabolic pathway introduced into these bacteria, we hypothesized that these engineered bacteria may utilize ornithine as a substrate to synthesize arginine, and thereby correcting the arginine: ornithine imbalance in the TME. To test this prediction, we assayed whether the arginine synthetic (Arg-ECN) could synthesize arginine from media devoid of ammonia but replete with ornithine. We found that Arg-ECN avidly synthesized arginine from the ornithine supplied in the media, even without exogenous ammonia (Fig. s6). Such findings indicate that this Arg-ECN strain could be used to tune the balance of arginine:ornithine in the TME to improve checkpoint inhibitor treatments and to potentiate anti-tumor immunity.

One of the unresolved questions from our work is why the physical properties of the TME would promote collagen-ECM synthetic TAM phenotype at the expense of anti-tumor inflammation? We hypothesize that the physical properties of the TME subvert anti-tumor responses in favor of pro-tumor ECM synthetic TAM programming due to an evolutionarily-beneficial, but maladaptive-in-the-tumor-context, wound-healing response triggered by the fibrillar ECM content of the TME or site of injury. Open and closed wounds initially display inflammatory programs regardless of sterility. These inflammatory programs are initiated by endogenous Damage Associated Molecular Patterns (DAMPs) in sterile wounds that collaborate with exogenous pattern associated molecular patterns (PAMPs) in open wounds to elicit inflammatory programs that remodel the site of injury for efficient surveillance and clearance of the sources of PAMPs/DAMPs. However, inflammatory programming is not static, as PAMPs/DAMPs are cleared, temporary reconstructive scaffolding (e.g., fibrin) is used to physically support the site of inflammation that is subsequently replaced by a more permanent fibrillar collagen matrix. This fibrillar collagen matrix is synthesized and mechanically entrained by neighboring cells (e.g., fibroblasts or keratinocytes), and it can dictate the recruitment of immune cells ^64^ or a programmatic homeostasis between stromal cell types ^79^. Similar to wound healing responses, the inflammatory programming of myeloid cells is not static. Dynamic phenotypic transitions occur in phases at sites of injury where myeloid-derived cells that initially expressed DAMP-driven pro-inflammatory signatures transition into anti-inflammatory states that produce TGFβ1 ^80^ and synthesize specific types of ECM that facilitate and typify the resolution of wound healing ^81^.

Physically, the similarities between the phased responses observed in wound healing or tumor progression are intuitively obvious. The early inflamed (i.e., PAMP/DAMP enriched) microenvironment is soft due to loss of tissue structure and cell death or clearance. As inflammatory signals are cleared or tolerated ^82^, ECM synthesis and cell contractility mediate tissue stiffening in specific regions of the TME (e.g., invasive borders). Microenvironmental physical cues synergize with PAMP/DAMP-signaling to inform infiltrating cells to express transcriptional phenotypes intended to match the stage of healing (e.g., fibronectin is a TLR4 ligand, that becomes inaccessible once incorporated into fibrillar matrix). Physical properties of healing wounds reflect the stage of repair, where a stiff wound has reached the phase in which the source of DAMP/PAMP has been cleared, and the wound is structurally entrained to its surrounding tissue. Our data suggest that one way that the physical properties of the TME drives TAMs to a pro-tumor/repair phenotype is by dictating the metabolic fate of arginine, which is a hallmark difference between pro/anti-inflammatory phenotypes of myeloid cells ^83^. Myeloid arginine metabolism serves an immunomodulatory role by restricting the proliferation and activity of T cells ^84^, which may serve to lessen inflammatory responses in healing wounds or progressing tumors ^85^. Overall, our results suggest that mechano-metabolic programming induced by the physical properties of a tissue may instruct and support infiltrating myeloid cell programming to match a presumed stage of wound healing, which is speciously subverted in tumor progression.

## Acknowledgments

**Funding:** This work was supported by 1F32CA236156-01A1, 5T32CA108462-15, and Sandler Program for Breakthrough Biomedical Research (postdoctoral Independence award) to K.M.T.; R35 CA242447-01A1, R01CA192914, and R01CA222508-01 to V.M.W.

**Author contributions:** Conceptualization: K.M.T. & V.M.W. Methodology: K.M.T, Investigation: K.M.T., K.K., O.M., G.T., C.S., J.t.H-S., F.C.P, I.B. and M-K.H. Formal Analysis: K.M.T., K.K., C.S., B.S., A.C., J.t.H-S., M-K.H., & A.J.I., Data curation: K.M.T. Funding acquisition: V.M.W., K.M.T, Project administration: K.M.T. Resources: Software: C.S. Supervision: V.M.W. Validation: K.M.T., K.K., G.T, and C.S. Visualization: K.M.T. Writing – original draft: K.M.T. Writing – review & editing: K.M.T., V.M.W., K.K., M-K.H., R.G., G.T., & A.C.

**Declaration of Interests:** The authors declare no competing interests.

## Methods

### Cell culture

Bone marrow was isolated from 8-12 week old female C57BL6/J femurs (via dissection and syringe-mediated PBS hydraulic pressure) and cultured in 25 mM glucose DMEM (Gen Clone, 25-500) supplemented with 10% Fetal Bovine Serum (FBS) (Hyclone, SH30071.03) and penicillin/streptomycin (Gibco, 151140-122) with 25 ng/mL recombinant murine Macrophage Colony Stimulating Factor (mCSF-1) (Peprotech, 315-02) for 7 days (media changes every 3 days, first 20 mL then 25 mL) in 15 cm^2^ tissue culture polystyrene culture plates (Gen Clone, 25-203) to generate bone marrow derived macrophages (BMDMs). BMDMs were treated with IL-4 [5 ng/mL] (Peprotech, 214-14) with and without TGFβ1 (R&D systems, 7666-MB).

CD8^+^ CTLs were isolated from the spleen, inguinal and axillary lymph nodes of 8-12 week old female C57BL6/J or OT1 mice (C57BL/6-Tg(TcraTcrb)1100Mjb/J, JAX:003831). Tissues were pulverized with a 1 mL syringe plunger through a PBS wetted 70 µm cell strainer (Corning, 352350). The resulting cell suspension was treated with ACK lysis buffer [150 mM NH_4_Cl, 10 mM KHCO_3_, 0.1 mM Na_2_EDTA, pH 7.3] to remove red blood cells and CD8^+^ T cells were isolated via EasySep Mouse CD8^+^ T Cell Isolation Kit (Stemcell Technologies,19853) per manufacturer’s instructions. CD8^+^ T cells were activated with Dynabeads Mouse T-Activator CD3/CD28 for T-Cell Expansion and Activation (Gibco, 11456D) without exogenous recombinant IL2 supplementation. OTI CD8^+^ T cells were isolated and co-cultured with PyMT-mCherryOVA tumor cells ^61^ that display the model antigen chicken ovalbumin (SIINFEKL) in 25 mM glucose DMEM (Gen Clone, 25-500) supplemented with or without L-ornithine (RPI, 020040).

### ECM coated polyacrylamide hydrogel cell culture surfaces (PA-gels)

Cleaned (10% ClO, 1M HCL, then 100% EtOH) round #1 German glass coverslips (Electron Microscopy Services) were coated with 0.5% v/v (3-Aminopropyl)triethoxysilane (APTES, Sigma, 440140), 99.2% v/v ethanol, and 0.3% v/v glacial acetic acid for 2 h and then cleaned in 100% EtOH on an orbital shaker at 22 °C. APTES activated coverslips were coated with PBS buffered acrylamide / bis-acrylamide (Bio-Rad, 1610140 and 1610142) solutions (3% / 0.05% for 400 Pa, 7.5% / 0.07% and 10% / 0.5% for 60k Pa) polymerized with TEMED (0.1% v/v) (Bio-Rad, 1610801) and Potassium Persulfate (0.1% w/v) (Fisher, BP180) to yield a final thickness of ∼ 85 µm. PA-gels were washed with 70% EtOH and sterile PBS prior 3,4-dihydroxy-L-phenylalanine (DOPA) coating for 5 min at 22 °C protected from light with sterile filtered DOPA in pH 10 [10 mM] Tris buffer ^86^. DOPA coated PA-gels were washed 2x with sterile PBS and ECM functionalized with 10 µg/mL Type I collagen (Corning, 354236) in sterile PBS 1 h at 37 °C.

### Fibroblast derived ECM culture surfaces

Fibroblast derived ECM (fECM) was generated based on previously described methods ^87^. Multipotent subcutaneous adipose derived fibroblasts ^88^ were seeded in gelatin-coated tissue culture dishes, cultured for 8 days in a complete medium (DMEM 10% FBS, 1% A/A) with 1 ng/mL TGFβ1 and 50 µg/ml ascorbic acid added every 48 h to support collagen synthesis. Cell cultures were washed with PBS and de-cellularized using a pre-warmed (37°C) extraction buffer for 2-5 min (0.5% Triton X-100, 20 mM NH4OH in PBS). Residual ECM were then washed three times with PBS prior to seeding BMDMs on top of the fECM.

### qPCR

Total RNA was isolated from biological samples with TRIzol (Invitrogen, 15596-018) according to the manufacturer’s instructions. cDNA was synthesized with 250 ng total RNA in 10 µL reaction volume with RNA using M-MLV reverse transcriptase (BioChain, Z5040002-100K) and 5X reaction buffer (BioChain, Z5040002-100K), random hexamers (Roche, 11034731001), dNTPs, and 1U of Ribolock (ThermoFisher, EO0384). RT-thermocycler program: random hexamers and RNA incubated at 70°C for 10 min, then held at 4°C until the addition of the M-MLV reverse transcriptase, dNTPs, Ribolock, and M-MLV-reverse transcriptase, then 50 °C for 1 h, 95 °C for 5 min, then stored at -20 °C until qPCR was performed. The reverse transcription reaction was then diluted to 50 µL total volume with ddH_2_O rendering a concentration of 20 ng RNA per 1 µL used in subsequent qPCR reactions. qPCR was performed in triplicate using PerfeCTa SYBR Green FastMix (Quantabio, Cat# 95072-05K) with an Eppendorf Mastercycler RealPlex^2^. qPCR thermocycler program: 95° C for 10 min, then 40 cycles of a 95°C for 15 s, 60°C for 20 s, followed by a melt curve 60-95°C over 10 min. Melt curves and gel electrophoresis were used to validate the quality of amplified products. The ΔCt values from independent experiments were used to calculate fold change of expression using the 2^-ΔΔCt^ method. For each gene measured, the SEM of the ΔCt values was calculated and used to generate positive and negative error values in the 2^-ΔΔCt^ fold change space. Plots of qPCR data display bars representing the mean fold change ±SEM and individual points representing the fold change value for each experiment relative to the mean.

### qPCR primers used

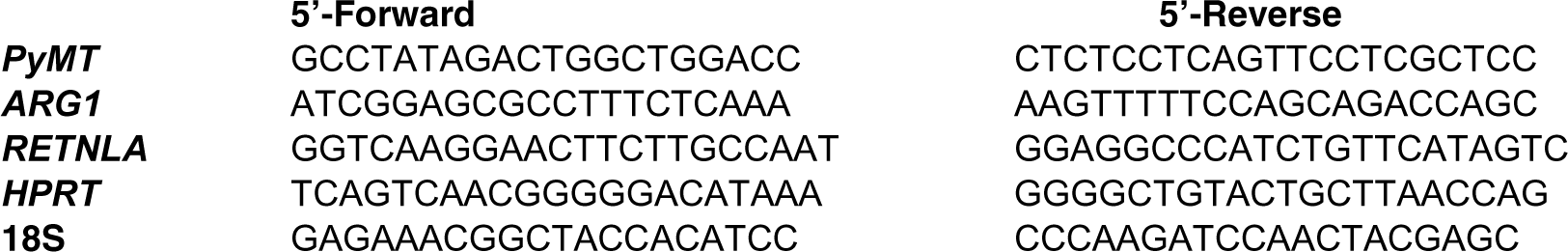

### In vitro respirometry

Mitochondrial stress tests were performed with a Seahorse XF24e cellular respirometer on non-permeablized cells (50k cells/well) in V7 microplates, with XF assay medium supplemented with 1 mM pyruvate (Gibco), 2 mM glutamine (Gibco),and 5 or 25 mM glucose (Sigma) at pH 7.4 and sequential additions via injection ports of Oligomycin [1 µM final], FCCP [1 µM final], and Antimycin A/Rotenone [1 µM final] during respirometry (concentrated stock solutions solubilized in 100% ethanol [2.5 mM] for mitochondrial stress test compounds). OCR values presented with non-mitochondrial oxygen consumption deducted.

### RNAseq

RNA-seq data from TAMs were generated with PyMT mammary tumors collected in DMEM on ice then minced with a prison shank-like razor blade. Tissues were digested in two 30 min rounds of collagenase and DNase with agitation at 37°C. At the end of each round, digested tissue was washed through a 100 µm cell strainer on ice. Red blood cells were lysed with ACK buffer [150 mM NH_4_Cl, 10 mM KHCO_3_, 0.1 mM Na_2_EDTA, pH 7.3] and remaining cells washed and counted. Cells were stained with anti-mouse CD45, anti-mouse CD11b, anti-mouse Ly6C, anti-mouse CD14, anti-mouse I-A/I-E, anti-mouse CD11c, anti-mouse CD24, anti-mouse CD29, anti-mouse CD31, and Zombie NIR live/dead. Sorted macrophages were CD45^+^, MHCII^+^, CD14^+^. Sorted cells were pelleted and snap frozen on dry ice. RNA was harvested using Trizol and measured on a spectrophotometer and run out on a Bioanalyzer to confirm quality. Libraries were prepared according to manufacturer instructions using Nugen Ovation plus Nextera. Multiplexed libraries were sequenced on an Illumina HiSeq4000, and reads were aligned to the human genome (hg19) using RNA STAR ^89^. Aligned reads were counted using HOMER ^90^, and hierarchical clustering was performed using Cluster ^91^ and visualized with Java TreeView. Gene Ontology and motif analysis was performed using Metascape ^92^. GEO Accession: GSE157290.

RNA-seq data from human tumors was re-analyzed for myeloid expression levels of transcripts associated with the collagen-containing ECM GO category, data published previously ^36^ (GEO Accession: GSE184398). Additionally, transcriptomic data for the BRCA, GBM, KIRC, LUAD, and PAAD projects were acquired from the TCGA repository. To control for the disparity in total RNA collected for each sample, observed library sizes were scaled into effective library sizes via the TMM method ^93^ in calcNormFactors (edgeR v3.34.1). Briefly, TMM assumes that most genes are not differentially expressed between samples and scales gene-level counts to alleviate artifactual expression differences that arise due to differing initial RNA abundances and sequencing depths. RNAseq counts were filtered to remove lowly expressed genes, then normalized using calcNormFactors (edgeR v3.34.1) and GSVA (v1.40.1) was applied to estimate enrichment of collagen genes as well as macrophage M1-and M2-like signatures driven by IL4 or LPS ^36^.

### Picrosirius red staining, polarized light microscopy, and second harmonic generation

Formalin Fixed Paraffin Embedded (FFPE) tissue sections were stained using 0.1% picrosirius red (Direct Red 80, Sigma, 365548) in picric acid solution (Sigma, P6744) and counterstained with Weigert’s hematoxylin (Cancer Diagnostics, catalogue number CM3951). Polarized light images were acquired using an Olympus IX81 microscope fitted with an analyzer (U-ANT) and a polarizer (U-POT, Olympus) oriented parallel and orthogonal to each other.

Second harmonic generation (SHG) imaging was performed using a custom-built two-photon microscope setup equipped resonant-scanning instruments based on published designs containing a five-photomultiplier tube (PMT) array (Hamamatsu, C7950). The setup consisted of a two-channel simultaneous video rate acquisition via two PMT detectors and an excitation laser (2□W MaiTai titanium-sapphire laser, 710–920□nm excitation range). SHG imaging was performed on a Prairie Technology Ultima System attached to an Olympus BX-51 fixed stage microscope equipped with a ×25 (numerical aperture 1.05) water immersion objective. Paraformaldehyde-fixed or FFPE tissue sections were exposed to polarized laser light at a wavelength of 830□nm and emitted light was separated using a filter set (short pass filter, 720□nm; dichroic mirror, 495□nm; band pass filter, 475/40□nm). Images of x–y planes at a resolution of 0.656Lmm per pixel were captured using at open-source Micro-Magellan software suite. Macrophages were depleted in MMTV-PyMT mice by intraperitoneal injections of 1 mg anti-CSF1 (Bio X Cell, catalogue number BE0204-A025, clone 5A1) or an IgG1 control (Bio X Cell, catalogue number BP0088-A025) every seven days starting at four weeks of age. Mice were sacrificed at eleven weeks of age for PS-red and polarized light microscopy.

### LC-MS metabolomics and arginine- or glutamine-derived ^13^C-flux

500k BMDMs were seeded on 50 mm^2^ varied stiffness ECM coated PA-gels and cultured for 24h in 25 mM glucose DMEM or 20-22 h in 25 mM glucose DMEM followed by medium change to DMEM lacking glutamine (Gibco, 11960044 replaced with with ^13^C_5_-glutamine (Cambridge Isotope Laboratories, CLM-1822-H-PK) for 2 h, or in SILAC DMEM medium (Gibco, 88364), lacking arginine and lysine, replaced with ^13^C_6_-arginine (Cambridge Isotope Laboratories, CLM-2265-H-PK) and L-lysine (sigma, L5501) for 4 h. 500k activated CD8^+^ T cells (CD3/CD28 Dynabeads) were cultured in DMEM supplemented with L-ornithine for prescribed time points with medium changed after 48 h for the 3 day activation. Cells were washed twice with PBS and extracted with mass spectrometry grade 80% methanol (Fisher, A456-1) and 20% water (Fisher, W6500) supplemented with 1 nmol DL-Norvaline (Sigma, N7502). Flash frozen tumors/tissues were pulverized with liquid nitrogen in a mortar and pestle on dry ice, 400 µg protein equivalents of tissue were extracted with mass spectrometry grade 80% methanol (Fisher, A456-1) and 20% water (Fisher, W6500) supplemented with 1 nmol DL-Norvaline (Sigma, N7502). Tissue or Insoluble material from cell extracts were used to calculate protein equivalents by resuspension in 0.2 M NaOH, heated to 95 °C for 25 min, and determined via BCA (Pierce, 23225). Dried metabolites were resuspended in 50% ACN:water and 1/10th of the volume was loaded onto a Luna 3 um NH2 100A (150 × 2.0 mm) column (Phenomenex). The chromatographic separation was performed on a Vanquish Flex (Thermo Scientific) with mobile phases A (5 mM NH_4_AcO pH 9.9) and B (ACN) and a flow rate of 200 µL/min. A linear gradient from 15% A to 95% A over 18 min was followed by 9 min isocratic flow at 95% A and reequilibration to 15% A. Metabolites were detected with a Thermo Scientific Q Exactive mass spectrometer run with polarity switching (+3.5 kV / -3.5 kV) in full scan mode with an m/z range of 65-975. TraceFinder 4.1 (Thermo Scientific) was used to quantify the targeted metabolites by area under the curve using expected retention time and accurate mass measurements (< 5 ppm). Values were normalized to cell number and sample protein concentration. Relative amounts of metabolites were calculated by summing up the values for all isotopologues of a given metabolite. Fractional contributional (FC) of ^13^C carbons to total carbon for each metabolite was calculated as previously described ^94^. Data analysis was accomplished with an in-house developed R scripts.

### Immunofluorescence microscopy

Cells or tissues were fixed in 4% paraformaldehyde (Electron Microscopy Services, 15710) in PBS for 30 min at room temperature for cell cultures or overnight at 4 °C for tissues, washed and blocked with a blocking buffer (HBSS fortified with: 10% FBS (Hyclone), 0.1% BSA (Fisher, BP1600), 0.05% saponin (EMD, L3771), and 0.1% Tween 20 (Fisher, BP337500). Primary antibodies [1:100-1:200] for 2 h at RT (22-23 °C) or 24 h at 4 °C, Secondary antibodies [1:1000] for 2 h at RT. Samples were imaged with a Nikon Eclipse Ti spinning disc microscope, Yokogawa CSU-X, Andor Zyla sCMOS, Andor Multi-Port Laser unit, and Molecular Devices MetaMorph imaging suite. Antibodies: Cleaved Caspase 3 (Cell Signaling, 9661), Collagen XII (Sigma, SAB4500395), Collagen VI (Abcam, ab6588), PyMT9 SCBT, PyMT (SCBT, sc-53481), and F4/80 (Cell Signaling, 70076).

### Fibrillar collagen tumor organoids and orthotopic tumors

Collagen type I [4.8 mg/mL final] (Rat-tail, Corning, 354249) was suspended in 0.1% glacial acetic acid with (stiff) or without (soft) non-metabolizable L-ribose [0.5 mM final] (Chem Impex International, 28127) for at least 28 days at 4 °C to facilitate complete collagen glycation/crosslinking ^32^. To generate orthotopic tumors or 3D-tumor cell organoids, 1 M/mL PyMT tumor cells were suspended in collagen +/-L-ribose supplemented with basement membrane extract [20 % v/v final (R&D Systems, Cultrex BME, 3532-005-02) and DMEM [1x final] and the pH was adjusted to ∼7.3. Tumor cell-collagen suspensions were kept on ice until implantation via 25 g syringe (100 µL per animal) or pipetted (20 µL) into culture dishes containing growth media for in vitro tumor organoid cultures.

### Primary tumor growth, circulating tumor cells, and lung metastasis

Tumor growth was monitored by caliper measurement and volume estimation based on V = (W2 × L)/2 or absolute measurement of mass. Lung metastases were quantified via immunofluorescence microscopy (PyMT) of lung sections.

All mouse studies were maintained under pathogen-free conditions and performed performed with strict adherence to the use authorizations (AN194983 and AN184232) provided by the University of California, San Francisco Institutional Animal Care and Use Committee (IACUC).

### Amino acid supplementation and PD-1 checkpoint blockade

Fibrillar collagen orthotopic tumors were generated with 100k PyMT cells in 100 µL collagen suspension per mouse, implanted via 25g syring injection into the 4/5 mammary fat pad of 8 week old female C57BL6/J mice. Every day until humane endpoint the mice were gavaged with 100 µL water or 2 g/kg glycine (Fisher, BP3815), ornithine (RPI, 020040), or arginine (TCI, A0528) suspended in water (∼100 µL). For PD-1 checkpoint blockade, mice were gavaged with glycine, ornithine, or arginine daily for the two weeks after orthotopic tumor implantation, and on day 5, 9, and 13 after orthotopic tumor implantatio n mice were intraperitoneally injected with 200 μg of anti-PD-L1 monoclonal antibody (RMP1-14, BioXCell, BE0146, Lot: 810421D1) or isotype control (IgG2a, BioXCell, BE0089, Lot: 815021S1) For PD-1 checkpoint blockade, mice were humanely euthanized when the tumors passed 1000 mm^3^ (no tumors exceeded 2 cm in diameter).

### Fluorescence activated cell sorting

FACS was performed as described previously ^95^. Tumors were processed into single cell suspensions via razor blade-mediated pulverization and enzymatic digestion with DNAse [200 mg/mL] (Sigma-Aldrich, 10104159001), Collagenase I [100 U/mL], and Collagenase Type IV [500 U/mL] (Worthington Biochemical, LS004197 and LS004189) on a shaking incubator (37°C) for 30 min. Enzymatic activity was quenched by doubling the volume of the tissue digest with FACS buffer (2% FCS in PBS). The resulting cell suspensions were filtered through 70 µm cell strainer (Corning, 352350) to obtain single cell suspensions. Lymph nodes and tumor-draining lymph nodes were pulverized with a 1 mL syringe plunger through a PBS wetted 70 µm cell strainer to generate single cell suspensions. For each sample, 5-10 x 10^6^ cells were washed with PBS and stained with Zombie NIR Fixable live/dead dye (Biolegend, 423106) at 4 °C for 20 min. Cells were washed with PBS followed by surface staining at 4 °C with directly conjugated antibodies (see list below) diluted in FACS buffer containing anti-CD16/32 (BioXCell, BE0307, RRID:AB_2736987 or BioLegend, 101320, RRID:AB_1574975) for 30 min. Cells were washed again with FACS buffer. For intracellular staining, cells were restimulated in RPMI (Gibco, 11875085) supplemented with 10% FCS (Benchmark, NC1643060), Pen/Strep/Glut (Gibco, 10378016), and b-mercaptoethanol [50 µM] (Gibco, 21985023) containing phorbol 12-myristate 12-acetate (PMA) [50 ng/mL] (Sigma, P8139), ionomycin [500 ng/mL] (Invitrogen, I24222), brefeldin A (BFA) [3 mg/mL] (Sigma, B7651) in 5% CO^2^ at 37 °C for 3-5 h. After surface staining, cells were fixed at 4 °C for 20 min and washed in permeabilization buffer from the Fix/Perm kit (BD Biosciences, 554714). Antibodies against intracellular targets were diluted in permeabilization buffer and cells were incubated at 4 °C for 30 min followed by another wash prior to assessment with BD LSR Fortessa SORP cytometer. All flow cytometry data was generated and analyzed in a blinded fashion.

| **Antibody target – fluorophore (clone)** | **Supplier** | **Catalogue #; RRID** |
| --- | --- | --- |
| anti-mouse Ly6C - BV711 (clone HK1.4) | Biolegend | Cat# 128037; RRID: AB_2562630 |
| anti-mouse F4/80 - BV510 (clone BM8) | Biolegend | Cat# 123135; RRID: AB_2562622 |
| anti-mouse CD24 - PECy7 (clone M1/69) | Biolegend | Cat# 101822; RRID: AB_756048 |
| anti-mouse CD11c - BV650 (clone N418) | Biolegend | Cat# 117339; RRID: AB_2562414 |
| anti-mouse CD11b - PerCp-Cy5.5 (clone ICRF44) | Biolegend | Cat# 301328; RRID: AB_10933428 |
| anti-mouse I-A/I-E - PE (clone AF6-88.5) | Biolegend | Cat# 107630; RRID: AB_2069376 |
| anti-mouse CD24 - BV421 (clone M1/69) | Biolegend | Cat# 101825; RRID: AB_10901159 |
| anti-mouse CD38 - BV711 (clone 90/CD38) | BD BioSciences | Cat# 740697; RRID: AB_2740381 |
| anti-mouse CD38 - FITC (clone 90/CD38) | BD BioSciences | Cat# 558813; RRID: AB_397126 |
| anti-mouse CD4 - BUV395 (clone RM4-5) | BD BioSciences | Cat# 563790; RRID: AB_2738426 |
| anti-mouse CD44 - BUV737 (clone IM7) | BD BioSciences | Cat# 564392; RRID: AB_2738785 |
| anti-mouse CD44 - AF700 (clone IM7) | Thermo Fisher | Cat# 56-0441-80; RRID:AB_494012 |
| anti-mouse CD45 AF647 (clone 30-F11) | Biolegend | Cat# 103124; RRID: AB_493533 |
| anti-mouse CD45 - BUV395 (clone 30-F11) | BD BioSciences | Cat# 564279; RRID: AB_2651134 |
| anti-mouse CD45 - BV421 (clone 30-F11) | Biolegend | Cat# 304032; RRID: AB_2561357 |
| anti-mouse CD45.1 - BUV395 (clone A20) | BD BioSciences | Cat# 565212; RRID: AB_2722493 |
| anti-mouse CD45.1 - PECy7 (clone A20) | Biolegend | Cat# 110730; RRID: AB_1134168 |
| anti-mouse CD5 - BV510 (clone 53-7.3) | Biolegend | Cat# 100627; RRID: AB_2563930 |
| anti-mouse CD69 - BV650 (clone H1.2F3) | Biolegend | Cat# 104541; RRID: AB_2616934 |
| anti-mouse CD8 - AF488 (clone 53-6.7) | Biolegend | Cat# 100723; RRID: AB_389304 |
| anti-mouse CD8 - PerCp-Cy5.5 (clone 53-6.7) | Biolegend | Cat# 100734; RRID: AB_2075238 |
| anti-mouse H2kB - PE (clone AF6-88.5) | Biolegend | Cat# 116507; RRID: AB_313734 |
| anti-mouse IFNy - PE (clone XMG1.2) | Thermo Fisher | Cat# 12-7311-81; RRID: AB_466192 |
| anti-mouse IFNy - PECy7 (clone XMG1.2) | Biolegend | Cat# 505826; RRID: AB_2295770 |
| anti-mouse Ki67 - PE eFluor610 (clone SolA15) | Thermo Fisher | Cat# 61-5698-82; RRID: AB_2574620 |
| anti-mouse Ly6G - BV785 (clone 1A8) | Biolegend | Cat# 127645; RRID: AB_2566317 |
| anti-mouse I-A/I-E - AF700 (clone M5/114.15.2) | Biolegend | Cat# 107622; RRID: AB_493727 |
| anti-mouse PD-1 - BV605 (clone 29F.1A12) | Biolegend | Cat# 135219; RRID: AB_11125371 |
| anti-mouse TCF1 - PE (clone S33-966) | BD BioSciences | Cat# 564217; RRID: AB_2687845 |
| anti-mouse TOX - APC (clone REA473) | Miltenyi Biotec | Cat# 130-118-335; RRID: AB_2751485 |

### Data availability

RNA-seq data described here has been deposited in GEO and is publicly available (Accession: GSE157290 and GSE184398), and data for the BRCA, GBM, KIRC, LUAD, and PAAD projects were acquired from the TCGA repository https://portal.gdc.cancer.gov/.

---

**Figure S1:**
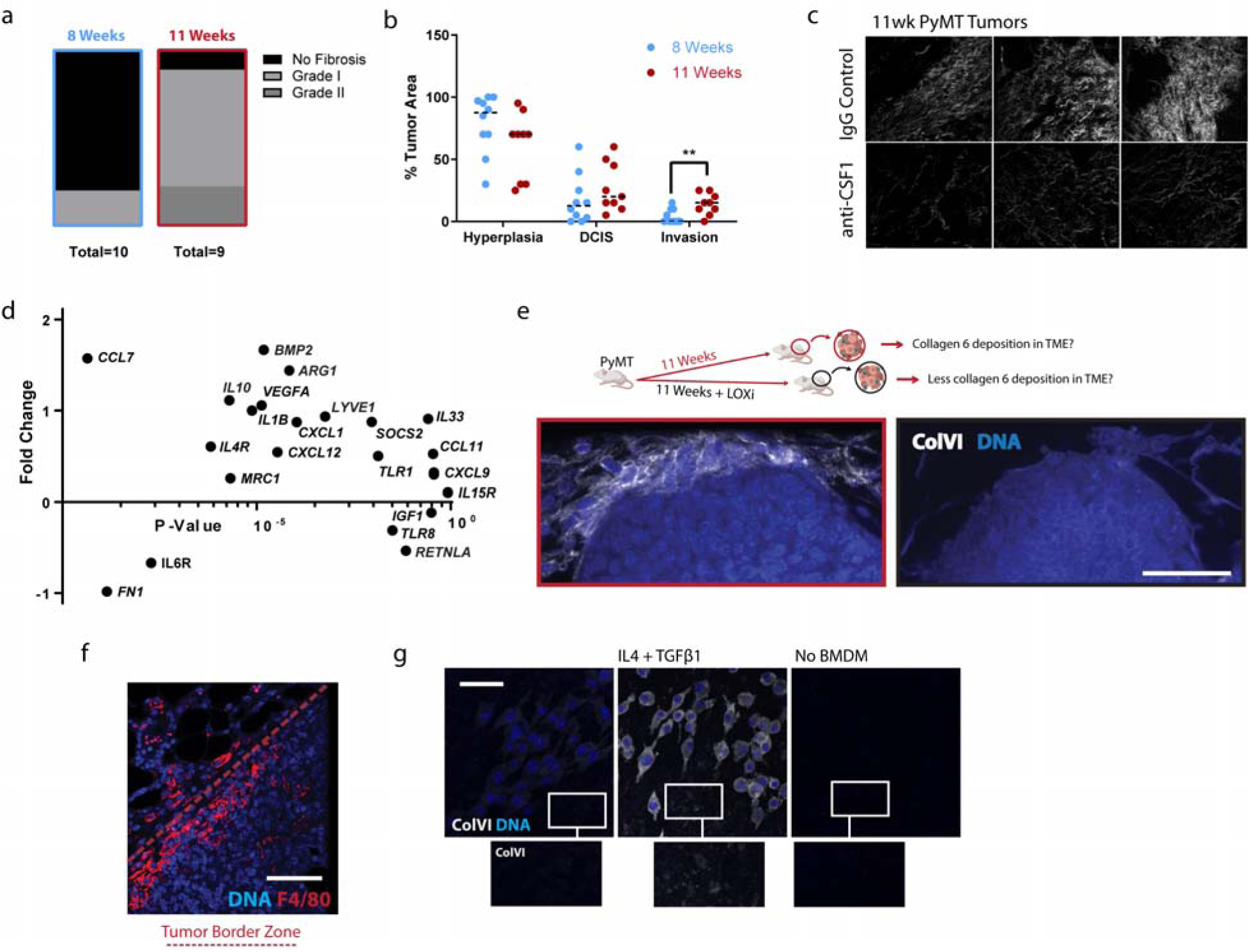
a. H&E-stained histological sections of 8 week and 11 week PyMT mammary tumors, quantification of the grade of stromal fibrosis. b. Quantification of pathological assessment for area of hyperplasia, DCIS with early invasion (DCIS) and advanced invasion (invasion) within H&E-stained histological sections of 8 week and 11 week PyMT mammary tumors (n=10 or 9). c. Representative second-harmonic generation (SHG) images of collagen fibers in 11 week old PyMT mammary tumors, treated with anti-CSF1 blocking antibody or IgG control weekly from 4 weeks of age until 11 weeks of age (n=3). d. Relative expression of macrophage polarization-associated gene expression of TAMs derived from 11 week old PyMT mammary tumors, relative to TAMs derived from 8 week old PyMT mammary tumors, (n=5). e. Representative immunofluorescence microscopy of Collagen VI (white) and DNA (blue) of 11 week old PyMT mammary tumors treated with or without LOX-inhibition, (Scale Bar: 100 µm). f. Representative immunofluorescence microscopy of F4/80 (red) and DNA (blue) in 11 week old PyMT mammary tumors (Scale Bar: 40 µm). g. Representative immunofluorescence microscopy of Collagen VI (white) and DNA (blue) of BMDMs treated with or without 5 ng/mL IL4 and 1 ng/mL TGFβ1 on fibroblast synthesized ECM surfaces for 24 h, (scale bar: 40 µm). *Data shown represent ± SEM. ***P < 0.005 via two-tailed unpaired Student t test (b)*

**Figure s2:**
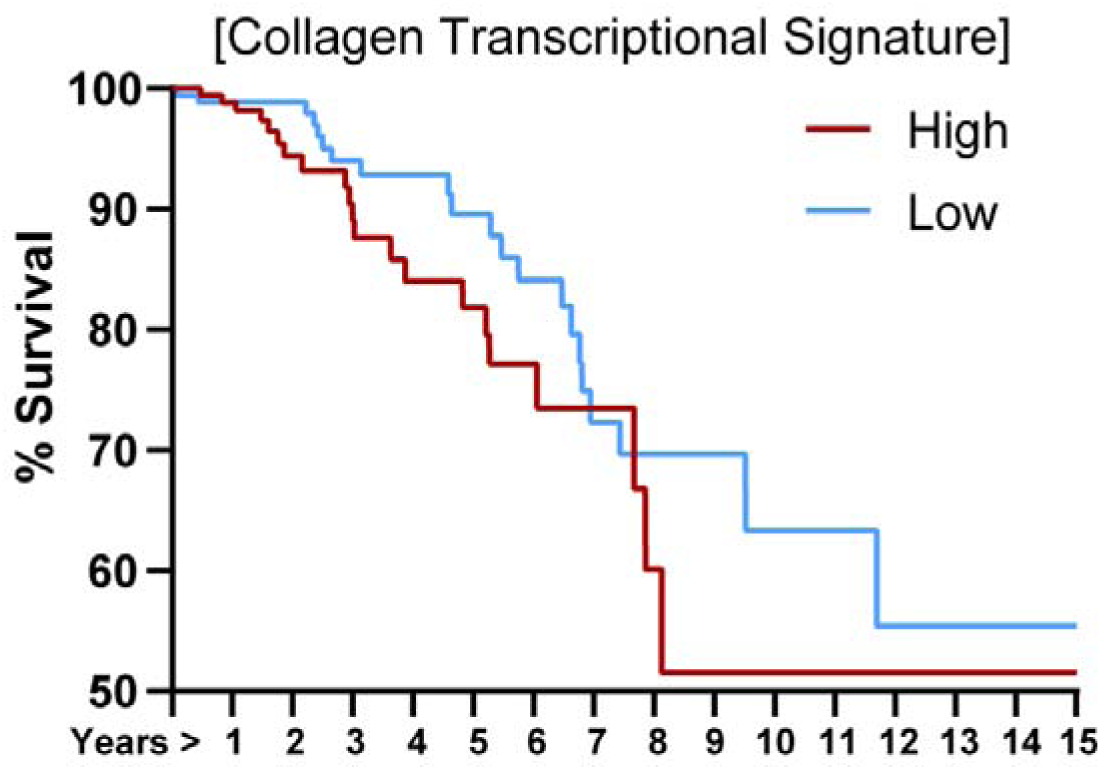
Kaplan-Meier survival curves of 2506 stage II and IIIA breast tumors, stratified for the top and bottom quartile expression level of the genes comprising the top GO category identified in d.

**Figure S3:**
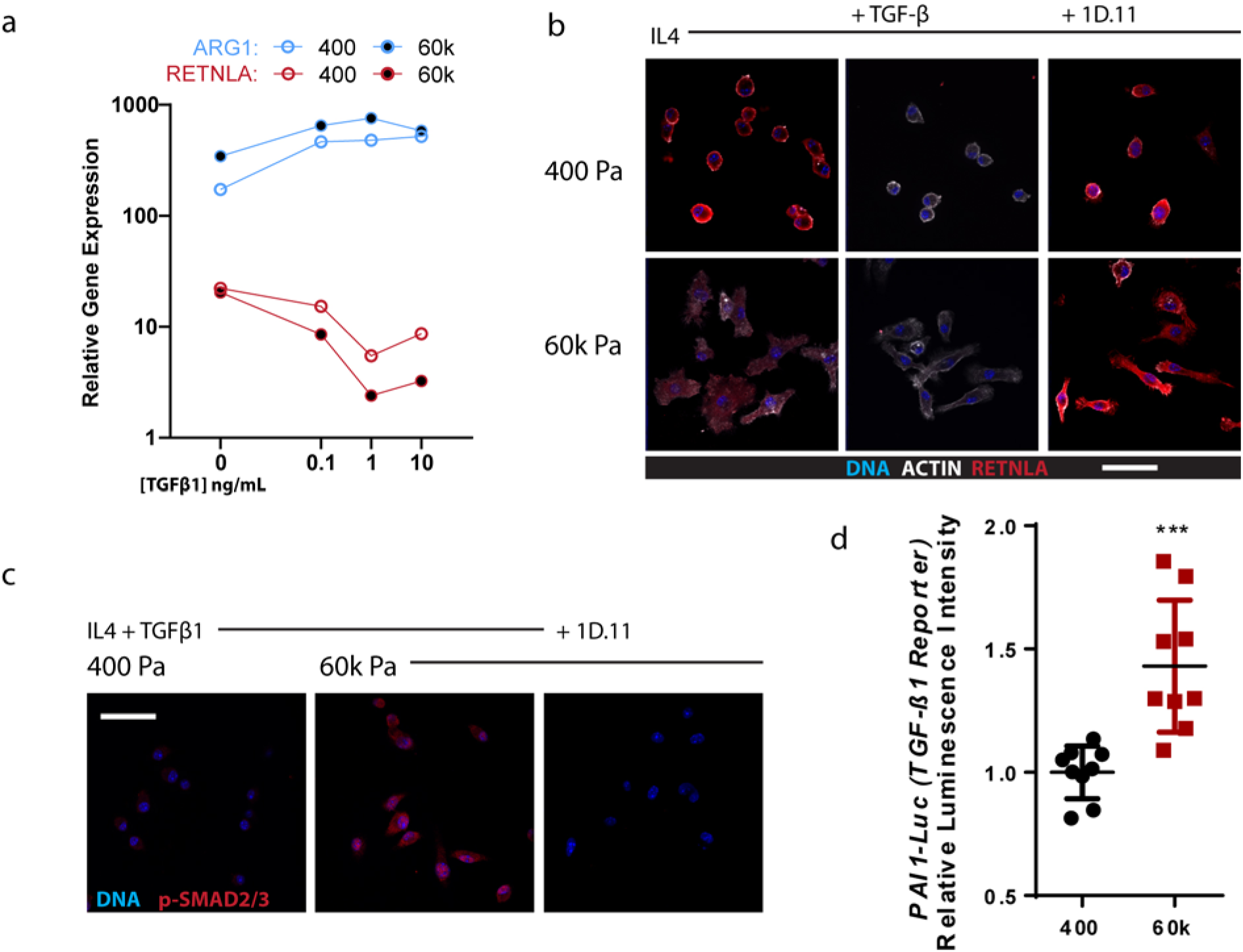
a. Relative gene expression of IL4-polarized BMDMs cultured on soft (400 Pa) or stiff (60k Pa) collagen I-coated polyacrylamide hydrogel surfaces treated with 0, 0.1, 1, or 10 ng/mL TGFβ1 for 4 h, qPCR-ΔΔCT normalized to BMDMs treated without IL4 (housekeeping gene: 18s), (n=3). b. Representative immunofluorescence microscopy for RETNLA (red) and DNA (blue) of IL4-polarized BMDMs cultured on soft (400 Pa) or stiff (60k Pa) collagen I-coated polyacrylamide hydrogel surfaces, treated with 1 ng/mL TGFβ1 with or without 10 µg/mL TGFβ1-blocking antibody (1D.11) for 24 h, (scale bar: 20 µm). d. Representative immunofluorescence microscopy for phosphorylated-SMAD2/3 (red) and DNA (blue) of IL4-polarized BMDMs cultured on soft (400 Pa) or stiff (60k Pa) collagen I-coated polyacrylamide hydrogel surfaces, treated with 1 ng/mL TGFβ1 with or without 10 µg/mL TGFβ1-blocking antibody (1D.11) for 24 h, (scale bar: 40 µm). e. Relative luminescent intensity of TGFβ1-reporter (PAI-1 Luc) expressing mink lung epithelial cells ^96^ when cultured for 24h in 1:1 conditioned media from IL4-polarized BMDMs cultured on soft (400 Pa) or stiff (60k Pa), (n=9).

**Figure S4:**
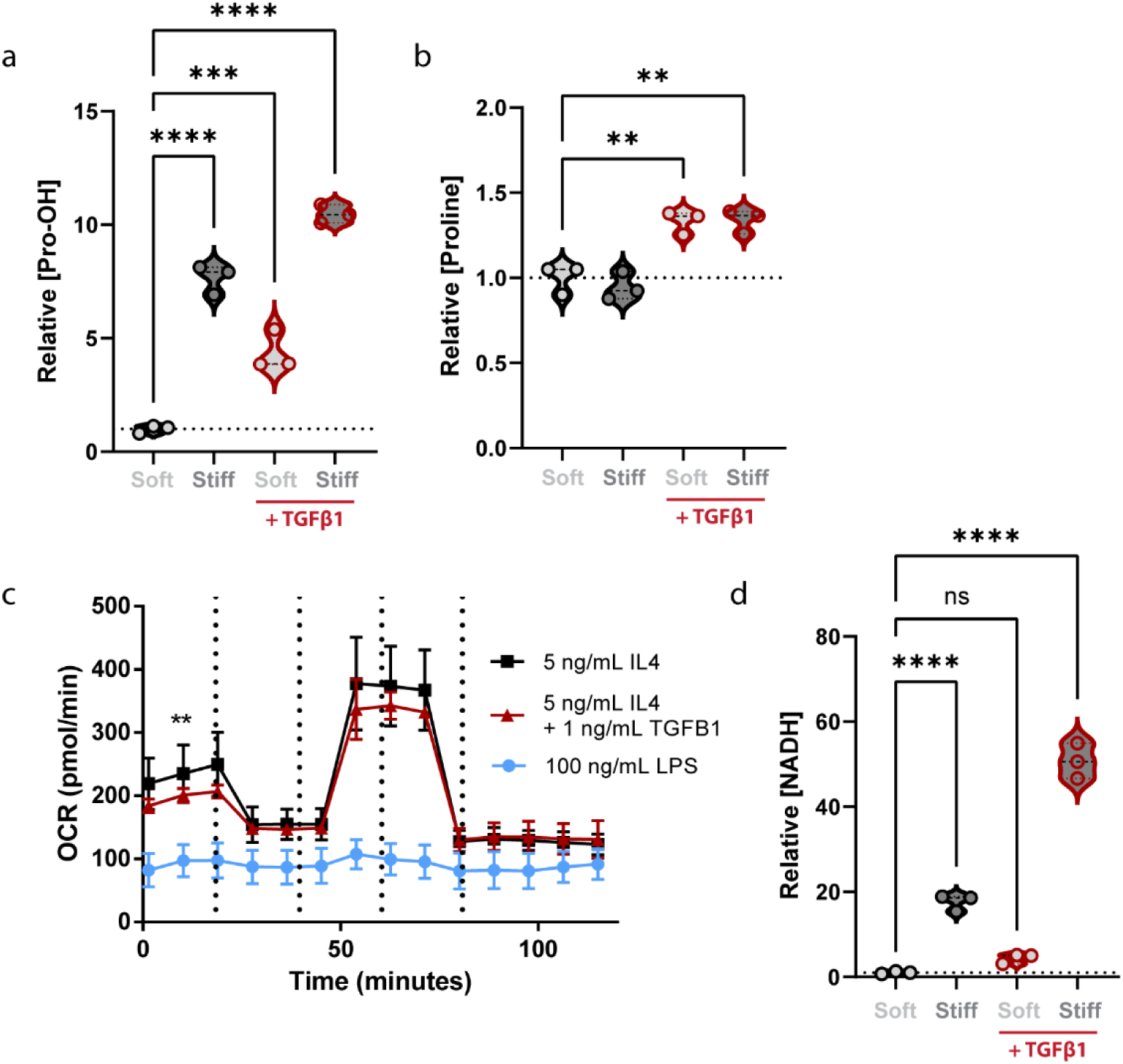
a. Relative hydroxyproline (Pro-OH) concentration in BMDMs cultured on soft (400 Pa) or stiff (60k Pa) collagen I-coated polyacrylamide hydrogel surfaces with or without 1 ng/mL TGFβ1 for 24 h, measured via LC-MS, (n=3). b. Relative proline concentration in BMDMs cultured on soft (400 Pa) or stiff (60k Pa) collagen I-coated polyacrylamide hydrogel surfaces with or without 1 ng/mL TGFβ1 for 24 h, measured via LC-MS, (n=3). c. Cellular respirometry of BMDMs cultured with or without 1 ng/mL TGFβ1 for 24h or 100 ng/mL LPS for 1 h along with sequential additions via injection ports of Oligomycin [1 µM final], FCCP [1 µM final], and Antimycin A/Rotenone [1 µM final] during respirometry measurements, measured with an SeahorseXF24, (n=5 wells, repeated 3 times) d. Relative NADH concentration in BMDMs cultured on soft (400 Pa) or stiff (60k Pa) collagen I-coated polyacrylamide hydrogel surfaces with or without 1 ng/mL TGFβ1 for 24 h, measured via LC-MS, (n=3). *Data shown represent ± SEM. **P < 0.01 or ****P < 0.0001 via one-way ANOVA with Tukey test for multiple comparisons (a, b, and d) and **P < 0.01 via two-tailed unpaired Student t test (c), ns indicates statistically not significant*.

**Figure S5:**
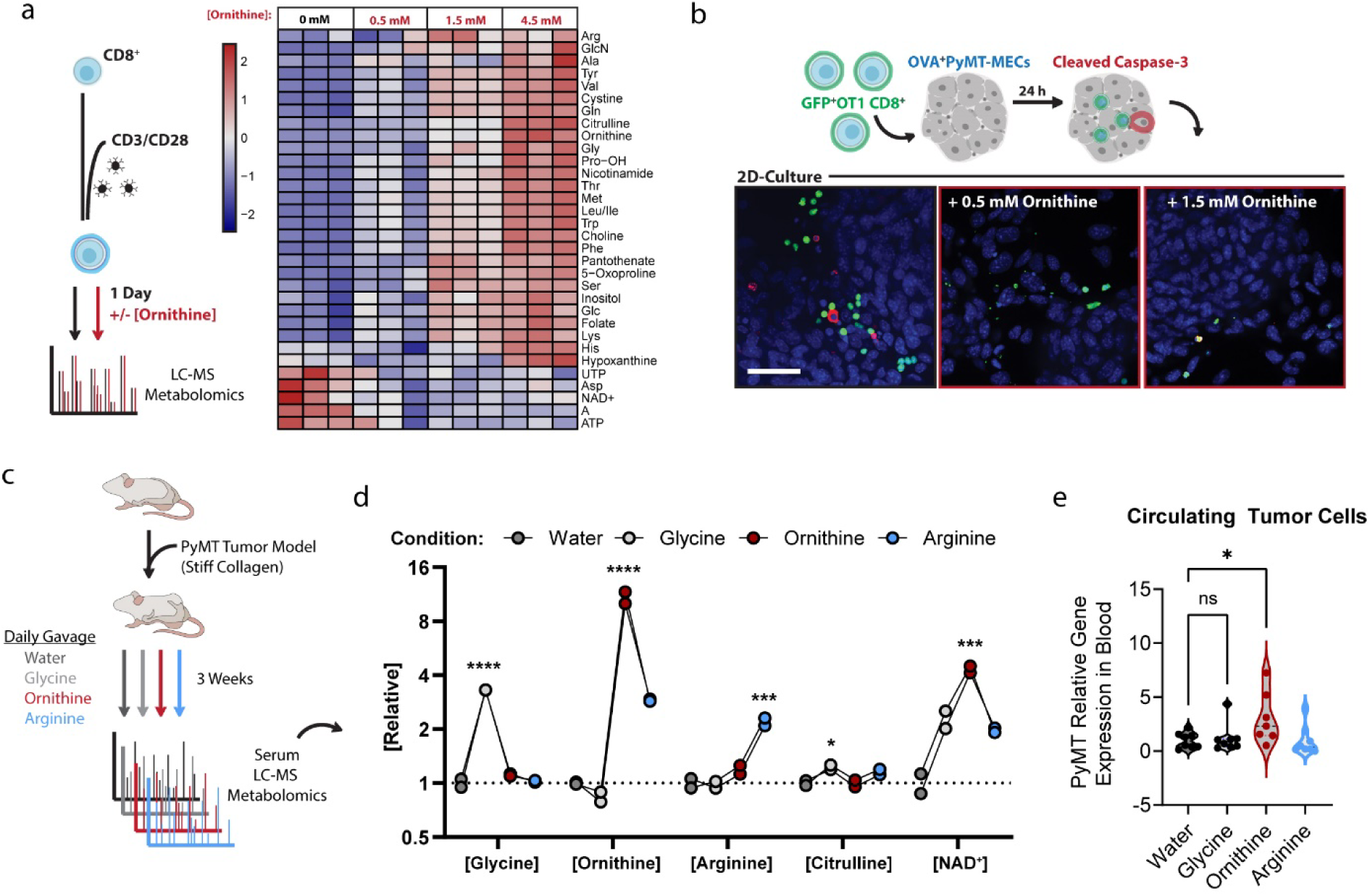
a. Heat map of relative metabolite levels of CD3/CD28-activated CD8^+^ CTLs cultured for 24 h in medium containing a molar ratio of 1:1 [0.5 mM], 3:1 [1.5 mM], and 9:1 [4.5 mM] ornithine:arginine, LC-MS analysis. (n=3 biological replicates) b. Representative immunofluorescence microscopy of OVA-PyMT tumor cells challenged with GFP^+^-OTI CTLs (green) for 24 h in medium containing a molar ratio of 1:1 [0.5mM] or 3:1 [1.5 mM] ornithine:arginine, cleaved-caspase 3 (red) and DNA (blue), (Scale Bar: 40 µm). c. Graphical description of the experimental setup for d-j. d. Relative serum metabolites derived from retro-orbital isolated blood from mice containing PyMT tumors after 3 weeks of growth in C57BL6/J mice gavaged daily with 2 g/kg glycine, ornithine, arginine, or water (100 µL), 4 h prior to isolation of blood/serum, measured with LC-MS (n=8 pooled into two technical replicates). e. qPCR based assessment of circulating tumor cells via PyMT expression in blood clot isolated from blood collected for serum metabolite analysis (e), qPCR-ΔΔCT (housekeeping gene: 18s), (n=8). *Data shown represent ± SEM. *P < 0.05, **P < 0.01, ***P < 0.005, or ****P < 0.001 via one-way ANOVA with Tukey test for multiple comparisons (d-j)*.

**Figure s6:**
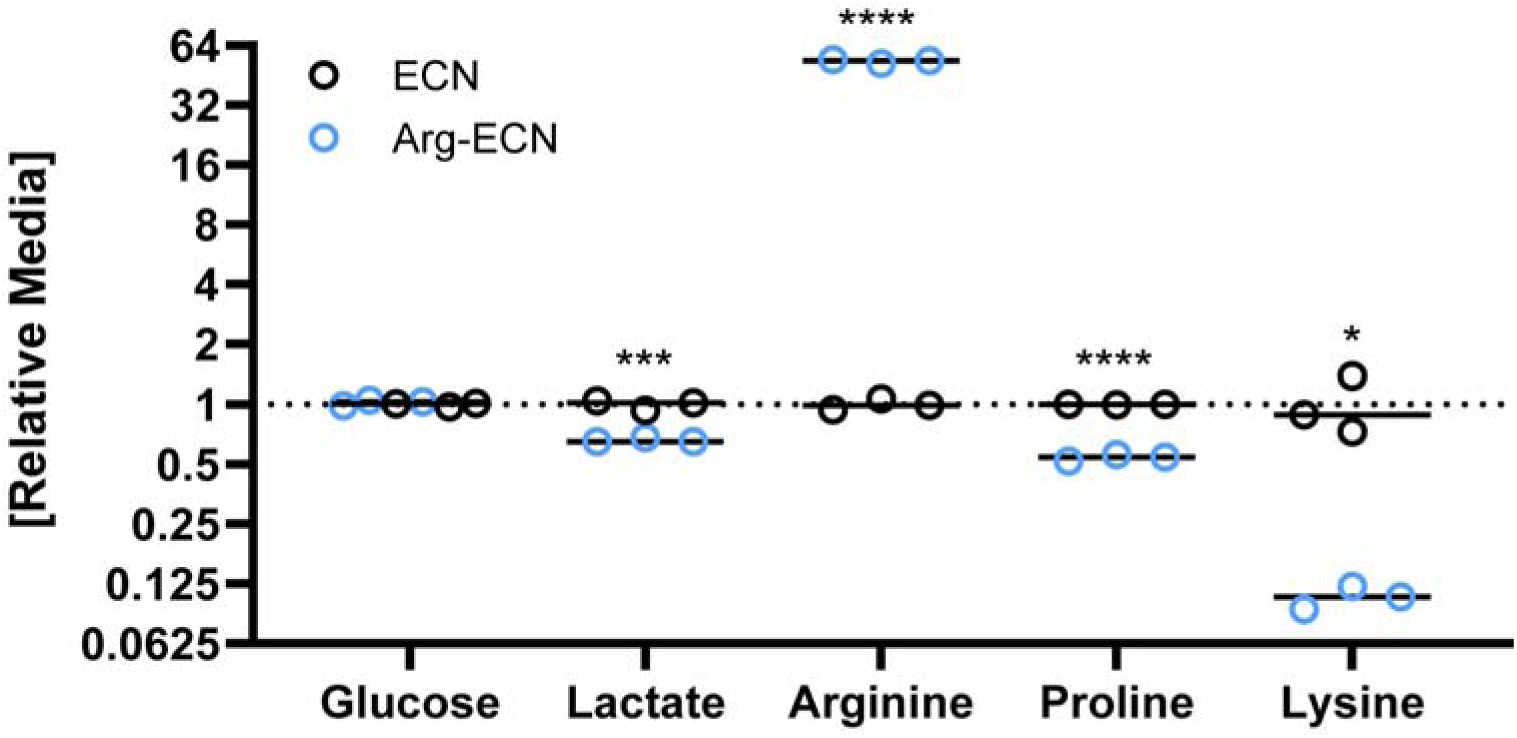
LC-MS based metabolomics of medium after 3 h of culture of Arg-ECN or ECN metabolizing a modified M9 medium, lacking ammonia, supplemented with ornithine [3 mM], (n=3).

## References

1. Galon, J. et al. Type, density, and location of immune cells within human colorectal tumors predict clinical outcome. Science (1979) 313, 1960–1964 (2006).

2. Tumeh, P. C. et al. PD-1 blockade induces responses by inhibiting adaptive immune resistance. Nature 2014 515:7528 515, 568–571 (2014).

3. Bagaev, A. et al. Conserved pan-cancer microenvironment subtypes predict response to immunotherapy. Cancer Cell 39, 845–865.e7 (2021).

4. Sun, X. et al. Tumour DDR1 promotes collagen fibre alignment to instigate immune exclusion. Nature 2021 599:7886 599, 673–678 (2021).

5. Nicolas-Boluda, A. et al. Tumor stiffening reversion through collagen crosslinking inhibition improves t cell migration and anti-pd-1 treatment. Elife 10, (2021).

6. Acerbi, I. et al. Human breast cancer invasion and aggression correlates with ECM stiffening and immune cell infiltration. Integrative Biology 7, 1120–1134 (2015).

7. Maller, O. et al. Tumour-associated macrophages drive stromal cell-dependent collagen crosslinking and stiffening to promote breast cancer aggression. Nature Materials 2020 20:4 20, 548–559 (2020).

8. Tauriello, D. V. F. et al. TGFβ drives immune evasion in genetically reconstituted colon cancer metastasis. Nature 2018 554:7693 554, 538–543 (2018).

9. Mariathasan, S. et al. TGFβ attenuates tumour response to PD-L1 blockade by contributing to exclusion of T cells. Nature 2018 554:7693 554, 544–548 (2018).

10. Chakravarthy, A., Khan, L., Bensler, N. P., Bose, P. & de Carvalho, D. D. TGF-β-associated extracellular matrix genes link cancer-associated fibroblasts to immune evasion and immunotherapy failure. Nature Communications 2018 9:1 9, 1–10 (2018).

11. Sahai, E. et al. A framework for advancing our understanding of cancer-associated fibroblasts. Nature Reviews Cancer 2020 20:3 20, 174–186 (2020).

12. Scharping, N. E. et al. Mitochondrial stress induced by continuous stimulation under hypoxia rapidly drives T cell exhaustion. Nature Immunology 2021 22:2 22, 205–215 (2021).

13. Buck, M. D., Sowell, R. T., Kaech, S. M. & Pearce, E. L. Metabolic Instruction of Immunity. Cell 169, 570–586 (2017).

14. Geiger, R. et al. L-Arginine Modulates T Cell Metabolism and Enhances Survival and Anti-tumor Activity. Cell 167, 829–842.e13 (2016).

15. Canale, F. P. et al. Metabolic modulation of tumours with engineered bacteria for immunotherapy. Nature 2021 598:7882 598, 662–666 (2021).

16. Ma, E. H. et al. Metabolic Profiling Using Stable Isotope Tracing Reveals Distinct Patterns of Glucose Utilization by Physiologically Activated CD8+ T Cells. Immunity 51, 856–870.e5 (2019).

17. Rodriguez, P. C. et al. l-Arginine Consumption by Macrophages Modulates the Expression of CD3ζ Chain in T Lymphocytes. The Journal of Immunology 171, 1232–1239 (2003).

18. Reinfeld, B. I. et al. Cell-programmed nutrient partitioning in the tumour microenvironment. Nature 2021 593:7858 593, 282–288 (2021).

19. Gabrilovich, D. I., Ostrand-Rosenberg, S. & Bronte, V. Coordinated regulation of myeloid cells by tumours. Nature Reviews Immunology 2012 12:4 12, 253–268 (2012).

20. Grzywa, T. M. et al. Myeloid Cell-Derived Arginase in Cancer Immune Response. Front Immunol 11, 938 (2020).

21. Wei, Z., Oh, J., Flavell, R. A. & Crawford, J. M. LACC1 bridges NOS2 and polyamine metabolism in inflammatory macrophages. Nature 2022 1–6 (2022) doi:10.1038/s41586-022-05111-3.

22. Menjivar, R. E. et al. Arginase 1 is a key driver of immune suppression in pancreatic cancer. bioRxiv 2022.06.21.497084 (2022) doi:10.1101/2022.06.21.497084.

23. Li, S. et al. Metabolism drives macrophage heterogeneity in the tumor microenvironment. Cell Rep 39, 110609 (2022).

24. Mouw, J. K. et al. Tissue mechanics modulate microRNA-dependent PTEN expression to regulate malignant progression. Nat Med 20, 360 (2014).

25. Taufalele, P. v., et al. Matrix stiffness enhances cancer-macrophage interactions and M2-like macrophage accumulation in the breast tumor microenvironment. Acta Biomater (2022) doi:10.1016/J.ACTBIO.2022.04.031.

26. Levental, K. R. et al. Matrix crosslinking forces tumor progression by enhancing integrin signaling. Cell 139, 891–906 (2009).

27. Afik, R. et al. Tumor macrophages are pivotal constructors of tumor collagenous matrix. Journal of Experimental Medicine 213, 2315–2331 (2016).

28. Schnoor, M. et al. Production of Type VI Collagen by Human Macrophages: A New Dimension in Macrophage Functional Heterogeneity. The Journal of Immunology 180, 5707–5719 (2008).

29. Biochemistry of Collagens, Laminins and Elastin. Biochemistry of Collagens, Laminins and Elastin (2016) doi:10.1016/C2015-0-05547-2.

30. Combes, A. J. et al. Discovering dominant tumor immune archetypes in a pan-cancer census. Cell 185, 184–203.e19 (2022).

31. Yu, X. et al. The Cytokine TGF-β Promotes the Development and Homeostasis of Alveolar Macrophages. Immunity 47, 903–912.e4 (2017).

32. Peranzoni, E. et al. Macrophages impede CD8 T cells from reaching tumor cells and limit the efficacy of anti–PD-1 treatment. Proc Natl Acad Sci U S A 115, E4041–E4050 (2018).

33. Sinha, P., Clements, V. K. & Ostrand-Rosenberg, S. Reduction of Myeloid-Derived Suppressor Cells and Induction of M1 Macrophages Facilitate the Rejection of Established Metastatic Disease. The Journal of Immunology 174, 636–645 (2005).

34. Leone, R. D. & Powell, J. D. Metabolism of immune cells in cancer. Nature Reviews Cancer 2020 20:9 20, 516–531 (2020).

35. Lim, A. R., Rathmell, W. K. & Rathmell, J. C. The tumor microenvironment as a metabolic barrier to effector T cells and immunotherapy. Elife 9, 1–13 (2020).

36. Bantug, G. R., Galluzzi, L., Kroemer, G. & Hess, C. The spectrum of T cell metabolism in health and disease. Nat Rev Immunol 18, 19–34 (2018).

37. Rossiter, N. J. et al. CRISPR screens in physiologic medium reveal conditionally essential genes in human cells. Cell Metab 33, 1248–1263.e9 (2021).

38. Leney-Greene, M. A., Boddapati, A. K., Su, H. C., Cantor, J. R. & Lenardo, M. J. Human Plasma-like Medium Improves T Lymphocyte Activation. iScience 23, 100759 (2020).

39. Cantor, J. R. et al. Physiologic Medium Rewires Cellular Metabolism and Reveals Uric Acid as an Endogenous Inhibitor of UMP Synthase. Cell 169, 258–272.e17 (2017).

40. Tharp, K. M. et al. Adhesion-mediated mechanosignaling forces mitohormesis. Cell Metab 33, 1322–1341.e13 (2021).

41. Sullivan, M. R. et al. Quantification of microenvironmental metabolites in murine cancers reveals determinants of tumor nutrient availability. Elife 8, (2019).

42. Kumar, V., Patel, S., Tcyganov, E. & Gabrilovich, D. I. The Nature of Myeloid-Derived Suppressor Cells in the Tumor Microenvironment. Trends Immunol 37, 208–220 (2016).

43. Raber, P., Ochoa, A. C. & Rodríguez, P. C. Metabolism of L-Arginine by Myeloid-Derived Suppressor Cells in Cancer: Mechanisms of T cell suppression and Therapeutic Perspectives. http://dx.doi.org/10.3109/08820139.2012.68063441, 614–634 (2012).

44. Caldwell, M. D., Mastrofrancesco, B., Shearer, J. & Bereiter, D. The temporal change in amino acid concentration within wound fluid--a putative rationale. Prog Clin Biol Res 365, 205–222 (1991).

45. Albaugh, V. L., Mukherjee, K. & Barbul, A. Proline Precursors and Collagen Synthesis: Biochemical Challenges of Nutrient Supplementation and Wound Healing. J Nutr 147, 2011–2017 (2017).

46. Mehl, A. A., Damião, A. O. M. C., Viana, S. D. D. O. & Andretta, C. P. Hard-to-heal wounds: A randomised trial of an oral proline-containing supplement to aid repair. J Wound Care 30, 26–31 (2021).

47. Schwörer, S. et al. Proline biosynthesis is a vent for TGFβ-induced mitochondrial redox stress. EMBO J 39, e103334 (2020).

48. Durante, W., Liao, L., Reyna, S. v., Peyton, K. J. & Schafer, A. I. Transforming Growth Factor-β1 Stimulates l-Arginine Transport and Metabolism in Vascular Smooth Muscle Cells. Circulation 103, 1121–1127 (2001).

49. Torrino, S. et al. Mechano-induced cell metabolism promotes microtubule glutamylation to force metastasis. Cell Metab 33, 1342–1357.e10 (2021).

50. Guo, L. et al. Kindlin-2 links mechano-environment to proline synthesis and tumor growth. Nature Communications 2019 10:1 10, 1–20 (2019).

51. Argüello, R. J. et al. SCENITH: A Flow Cytometry-Based Method to Functionally Profile Energy Metabolism with Single-Cell Resolution. Cell Metab 32, 1063–1075.e7 (2020).

52. Mak, T. W. et al. Glutathione Primes T Cell Metabolism for Inflammation. Immunity 46, 675–689 (2017).

53. Yarosz, E. L. & Chang, C. H. The Role of Reactive Oxygen Species in Regulating T Cell-mediated Immunity and Disease. Immune Netw 18, (2018).

54. Engelhardt, J. J. et al. Marginating Dendritic Cells of the Tumor Microenvironment Cross-Present Tumor Antigens and Stably Engage Tumor-Specific T Cells. Cancer Cell 21, 402– 417 (2012).

55. Zaitsev, A. et al. Precise reconstruction of the TME using bulk RNA-seq and a machine learning algorithm trained on artificial transcriptomes. Cancer Cell 40, 879–894.e16 (2022).

56. Joyce, J. A. & Fearon, D. T. T cell exclusion, immune privilege, and the tumor microenvironment. Science (1979) 348, 74–80 (2015).

57. Zhu, Y. et al. CSF1/CSF1R Blockade Reprograms Tumor-Infiltrating Macrophages and Improves Response to T Cell Checkpoint Immunotherapy in Pancreatic Cancer Models. Cancer Res 74, 5057 (2014).

58. Cannarile, M. A. et al. Colony-stimulating factor 1 receptor (CSF1R) inhibitors in cancer therapy. J Immunother Cancer 5, 1–13 (2017).

59. Laviron, M. et al. Tumor-associated macrophage heterogeneity is driven by tissue territories in breast cancer. Cell Rep 39, 110865 (2022).

60. Mujal, A. M. et al. Holistic Characterization of Tumor Monocyte-to-Macrophage Differentiation Integrates Distinct Immune Phenotypes in Kidney Cancer. Cancer Immunol Res 10, 403–419 (2022).

61. Cassetta, L. et al. Human Tumor-Associated Macrophage and Monocyte Transcriptional Landscapes Reveal Cancer-Specific Reprogramming, Biomarkers, and Therapeutic Targets. Cancer Cell 35, 588 (2019).

62. Jiang, H., Hegde, S. & DeNardo, D. G. Tumor-associated fibrosis as a regulator of tumor immunity and response to immunotherapy. Cancer Immunol Immunother 66, 1037–1048 (2017).

63. Tran, D. H. et al. Mitochondrial NADP+ is essential for proline biosynthesis during cell growth. Nature Metabolism 2021 3:4 3, 571–585 (2021).

64. Pakshir, P. et al. Dynamic fibroblast contractions attract remote macrophages in fibrillar collagen matrix. Nat Commun 10, (2019).

65. Pickup, M. W., Mouw, J. K. & Weaver, V. M. The extracellular matrix modulates the hallmarks of cancer. EMBO Rep 15, 1243–1253 (2014).

66. Ricard-Blum, S. The Collagen Family. Cold Spring Harb Perspect Biol 3, 1–19 (2011).

67. Papanicolaou, M. et al. Temporal profiling of the breast tumour microenvironment reveals collagen XII as a driver of metastasis. Nature Communications 2022 13:1 13, 1–21 (2022).

68. Simões, F. C. et al. Macrophages directly contribute collagen to scar formation during zebrafish heart regeneration and mouse heart repair. Nature Communications 2020 11:1 11, 1–17 (2020).

69. Wishart, A. L. et al. Decellularized extracellular matrix scaffolds identify full-length collagen VI as a driver of breast cancer cell invasion in obesity and metastasis. Sci Adv 6, (2020).

70. Thompson, S. B. et al. Formin-like 1 mediates effector t cell trafficking to inflammatory sites to enable t cell-mediated autoimmunity. Elife 9, 1–27 (2020).

71. Davidson, M. D., Burdick, J. A. & Wells, R. G. Engineered biomaterial platforms to study fibrosis. Adv Healthc Mater 9, e1901682 (2020).

72. Carey, S. P., Martin, K. E. & Reinhart-King, C. A. Three-dimensional collagen matrix induces a mechanosensitive invasive epithelial phenotype. Scientific Reports 2017 7:1 7, 1–14 (2017).

73. Riedel, S. et al. Design of biomimetic collagen matrices by reagent-free electron beam induced crosslinking: Structure-property relationships and cellular response. Mater Des 168, 107606 (2019).

74. Özdemir, B. C. et al. Depletion of carcinoma-associated fibroblasts and fibrosis induces immunosuppression and accelerates pancreas cancer with reduced survival. Cancer Cell 25, 719–734 (2014).

75. Germano, G. et al. Role of macrophage targeting in the antitumor activity of trabectedin. Cancer Cell 23, 249–262 (2013).

76. Mantovani, A., Allavena, P., Marchesi, F. & Garlanda, C. Macrophages as tools and targets in cancer therapy. Nature Reviews Drug Discovery 2022 1–22 (2022) doi:10.1038/s41573-022-00520-5.

77. Dröge, W. et al. Suppression of cytotoxic T lymphocyte activation by L-ornithine. The Journal of Immunology 134, (1985).

78. Lercher, A. et al. Type I Interferon Signaling Disrupts the Hepatic Urea Cycle and Alters Systemic Metabolism to Suppress T Cell Function. Immunity 51, 1074 (2019).

79. Adler, M. et al. Principles of Cell Circuits for Tissue Repair and Fibrosis. iScience 23, 100841 (2020).

80. Nguyen-Chi, M. et al. Identification of polarized macrophage subsets in zebrafish. Elife 4, (2015).

81. Foster, D. S., Jones, R. E., Ransom, R. C., Longaker, M. T. & Norton, J. A. The evolving relationship of wound healing and tumor stroma. JCI Insight 3, (2018).

82. Timblin, G. A. et al. Mitohormesis reprogrammes macrophage metabolism to enforce tolerance. Nature Metabolism 2021 3:5 3, 618–635 (2021).

83. Mills, C. D., Kincaid, K., Alt, J. M., Heilman, M. J. & Hill, A. M. M-1/M-2 Macrophages and the Th1/Th2 Paradigm. The Journal of Immunology 164, 6166–6173 (2000).

84. van de Velde, L. A. et al. T Cells Encountering Myeloid Cells Programmed for Amino Acid-dependent Immunosuppression Use Rictor/mTORC2 Protein for Proliferative Checkpoint Decisions *. Journal of Biological Chemistry 292, 15–30 (2017).

85. Eming, S. A., Murray, P. J. & Pearce, E. J. Metabolic orchestration of the wound healing response. Cell Metab 33, 1726–1743 (2021).

86. Wouters, O. Y., Ploeger, D. T. A., van Putten, S. M. & Bank, R. A. 3,4-Dihydroxy-L-Phenylalanine as a Novel Covalent Linker of Extracellular Matrix Proteins to Polyacrylamide Hydrogels with a Tunable Stiffness. Tissue Eng Part C Methods 22, 91–101 (2016).

87. Kaukonen, R., Jacquemet, G., Hamidi, H. & Ivaska, J. Cell-derived matrices for studying cell proliferation and directional migration in a complex 3D microenvironment. Nature Protocols 2017 12:11 12, 2376–2390 (2017).

88. Tharp, K. M. et al. Actomyosin-Mediated Tension Orchestrates Uncoupled Respiration in Adipose Tissues. Cell Metab 27, 602–615.e4 (2018).

89. Dobin, A. et al. STAR: ultrafast universal RNA-seq aligner. Bioinformatics 29, 15–21 (2013).

90. Lin, Y. C. et al. A global network of transcription factors, involving E2A, EBF1 and Foxo1, that orchestrates B cell fate. Nat Immunol 11, 635–643 (2010).

91. Eisen, M. B., Spellman, P. T., Brown, P. O. & Botstein, D. Cluster analysis and display of genome-wide expression patterns. Proc Natl Acad Sci U S A 95, 14863–14868 (1998).

92. Zhou, Y. et al. Metascape provides a biologist-oriented resource for the analysis of systems-level datasets. Nat Commun 10, 1523 (2019).

93. Robinson, M. D. & Oshlack, A. A scaling normalization method for differential expression analysis of RNA-seq data. Genome Biol 11, 1–9 (2010).

94. Cameron, A. M. et al. Inflammatory macrophage dependence on NAD(+) salvage is a consequence of reactive oxygen species-mediated DNA damage. Nat Immunol 20, 420– 432 (2019).

95. Kersten, K. et al. Spatiotemporal co-dependency between macrophages and exhausted CD8+ T cells in cancer. Cancer Cell 40, 624–638.e9 (2022).

96. Abe, M. et al. An assay for transforming growth factor-beta using cells transfected with a plasminogen activator inhibitor-1 promoter-luciferase construct. Anal Biochem 216, 276–284 (1994).

